# Distinct allosteric networks in CDK4 and CDK6 in the cell cycle and in drug resistance

**DOI:** 10.1101/2025.02.28.640857

**Authors:** Wengang Zhang, Devin Bradburn, Gretchen Heidebrink, Yonglan Liu, Hyunbum Jang, Ruth Nussinov, Mardo Kõivomägi

**Author notes:** Author for correspondence: Ruth Nussinov, Mardo Kõivomägi **Email:**.

## Abstract

Cyclin-dependent kinases 4 and 6 (CDK4 and CDK6) are key regulators of the G1-S phase transition in the cell cycle. In cancer cells, CDK6 overexpression often outcompetes CDK4 in driving cell cycle progression, contributing to resistance against CDK4/6 inhibitors (CDK4/6i). This suggests distinct functional and conformational differences between these two kinases, despite their striking structural and sequence similarities. Understanding the mechanisms that differentiate CDK4 and CDK6 is crucial, as resistance to CDK4/6i—frequently linked to CDK6 overexpression—remains a significant therapeutic challenge. Notably, CDK6 is often upregulated in CDK4/6i-resistant cancers and rapidly proliferating hematopoietic stem cells, underscoring its unique regulatory roles. We hypothesize that their distinct conformational dynamics explain their differences in phosphorylation of retinoblastoma protein, Rb, inhibitor efficacy, and cell cycle control. This leads us to question *how their dissimilar conformational dynamics encode their distinct actions*. To elucidate their differential activities, molecular mechanisms, and inhibitor binding, we combine biochemical assays and molecular dynamics (MD) simulations. We discover that CDK4 and CDK6 have distinct allosteric networks connecting the β3-αC loop and the G-loop. CDK6 exhibits stronger coupling and shorter path lengths between these regions, resulting in higher kinase activity upon cyclin binding and impacting inhibitor specificity. We also discover an unrecognized role of the unstructured CDK6 C-terminus, which allosterically connects and stabilizes the R-spine, facilitating slightly higher activity. Our findings bridge the gap between the structural similarity and functional divergence of CDK4 and CDK6, advancing the understanding of kinase regulation in cancer biology.

**Graphical abstract:** 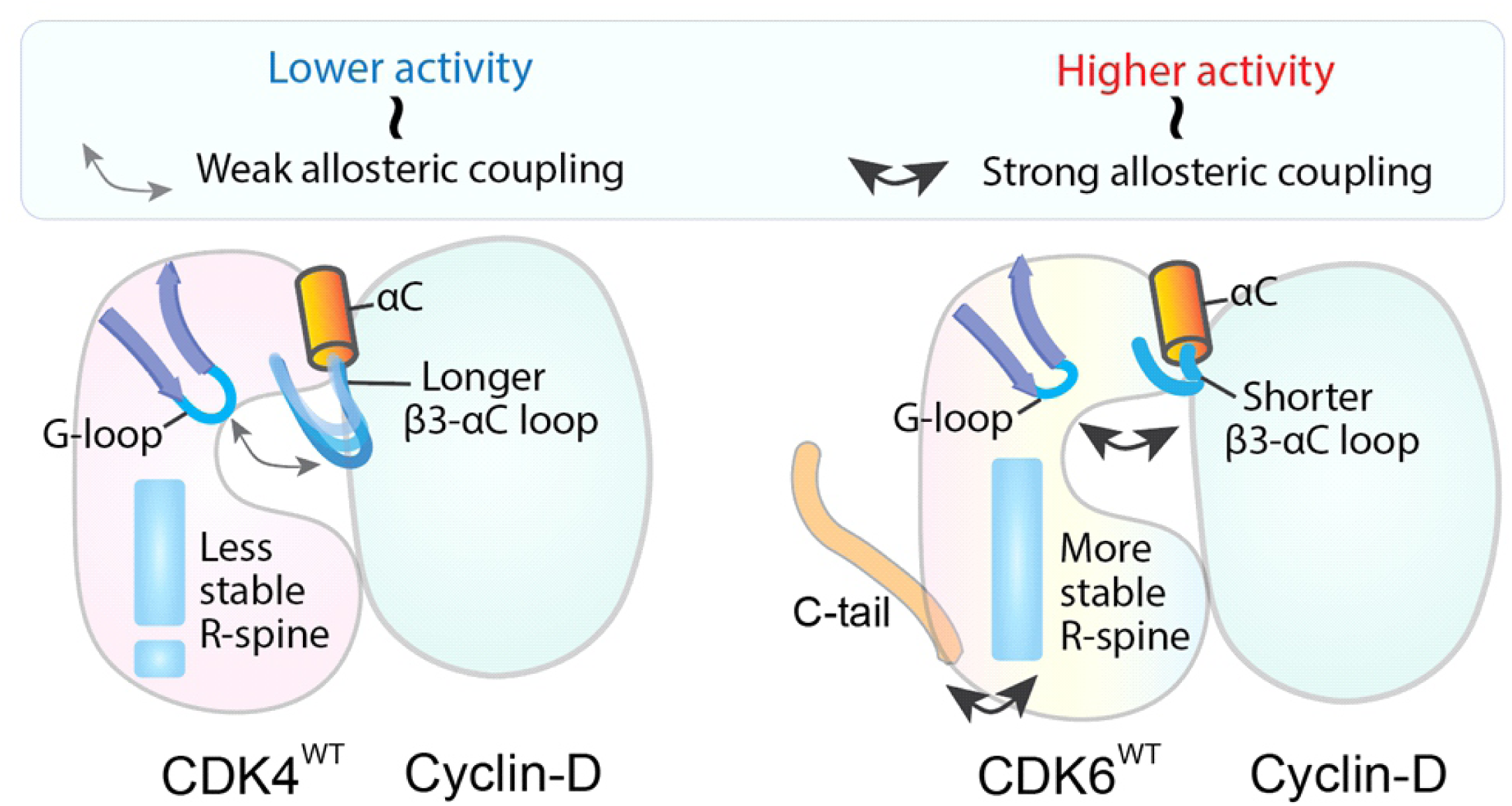

## 1 Introduction

The almost synonymous and interchangeable cyclin-dependent kinases 4 and 6 (CDK4 and CDK6), with relatively minor sequence and structural differences, differentially regulate the G_1_-S phase transition of the cell cycle in CDK4/6 inhibitor (CDK4/6i)-resistant cancer cells, adult tissues, and hematopoietic stem cells (HSCs).[1–5] The crucial question of the molecular mechanisms by which overexpressed CDK6 in cancer resistant cells facilitate cell cycle progression— in lieu of CDK4— remains unanswered.

Cyclin dependent kinases (CDKs) and their regulatory subunits, cyclins, are essential in the ordered progression of cells through the cell cycle.[6] Dysregulation of cyclin-CDK complexes leads to uncontrolled cell growth, making them promising targets for anticancer therapies.[7] Closely evolutionarily related, CDK4 and CDK6 are activated by cyclin D binding and phosphorylation by CDK7 **(Fig. S1)**.[8, 9] The two share overlapping functions in regulating progression through G_1_ by phosphorylating retinoblastoma protein (Rb).[10] Phosphorylation relieves Rb-mediated inhibition of E2F transcription factors, in turn stimulating expression of genes required for the transition into S-phase (such as cyclin E) and committing cells to replication **(Fig. 1A)**.[11] Because of their critical role in initiating the cell cycle, dual CDK4/6 inhibitors (palbociclib, abemaciclib, ribociclib) are used as chemotherapeutics in HR positive and HER2 negative breast cancer. However, their use is associated with a range of side effects, most notably neutropenia.[7, 12, 13] Hematopoietic stem cells (HSCs) rely on CDK6 to drive G_1_ progression, whereas most cell types, including breast cancer cells, utilize CDK4. Despite their sequence and structural similarity, CDK4 and CDK6 differentially regulate G_1_ progression in a variety of cell types and exhibit remarkable diversity in regulation, activation, and substrate recognition. In the case of rapidly dividing HSCs, the greater intrinsic catalytic activity of CDK6 may facilitate faster entry into the cell cycle. A CDK4 specific inhibitor, atirmociclib, has successfully been shown to halt breast cancer growth with reduced effect on neutrophil levels. However, an in depth understanding of the mechanistic differences between CDK4 and CDK6, which could inform better drug design, has not yet been established. This is likely in part due to existing static structural analyses failing to capture allosteric networks which modulate activity and substrate specificity through dynamic communication.[14] CDK4 and CDK6 likely rely on distinct allosteric pathways to mediate their unique functions, which may explain their specialized roles in the cell cycle and underlie differences in drug sensitivity.[15–19] These subtle structural differences may also underlie the development of drug resistance specifically through CDK6 overexpression. When in complex with the canonically inhibitory protein, INK4, CDK6 adopts a drug resistant yet ATP-binding competent state through which it can still sustain G_1_ progression.[38] The differential response to inhibitors suggests distinct allosteric mechanisms of CDK4 and CDK6 which are not yet understood.[3, 7, 20–25] Understanding these allosteric networks could be key to uncovering the unique mechanisms of these kinases and improved inhibitors.

**Figure 1.**
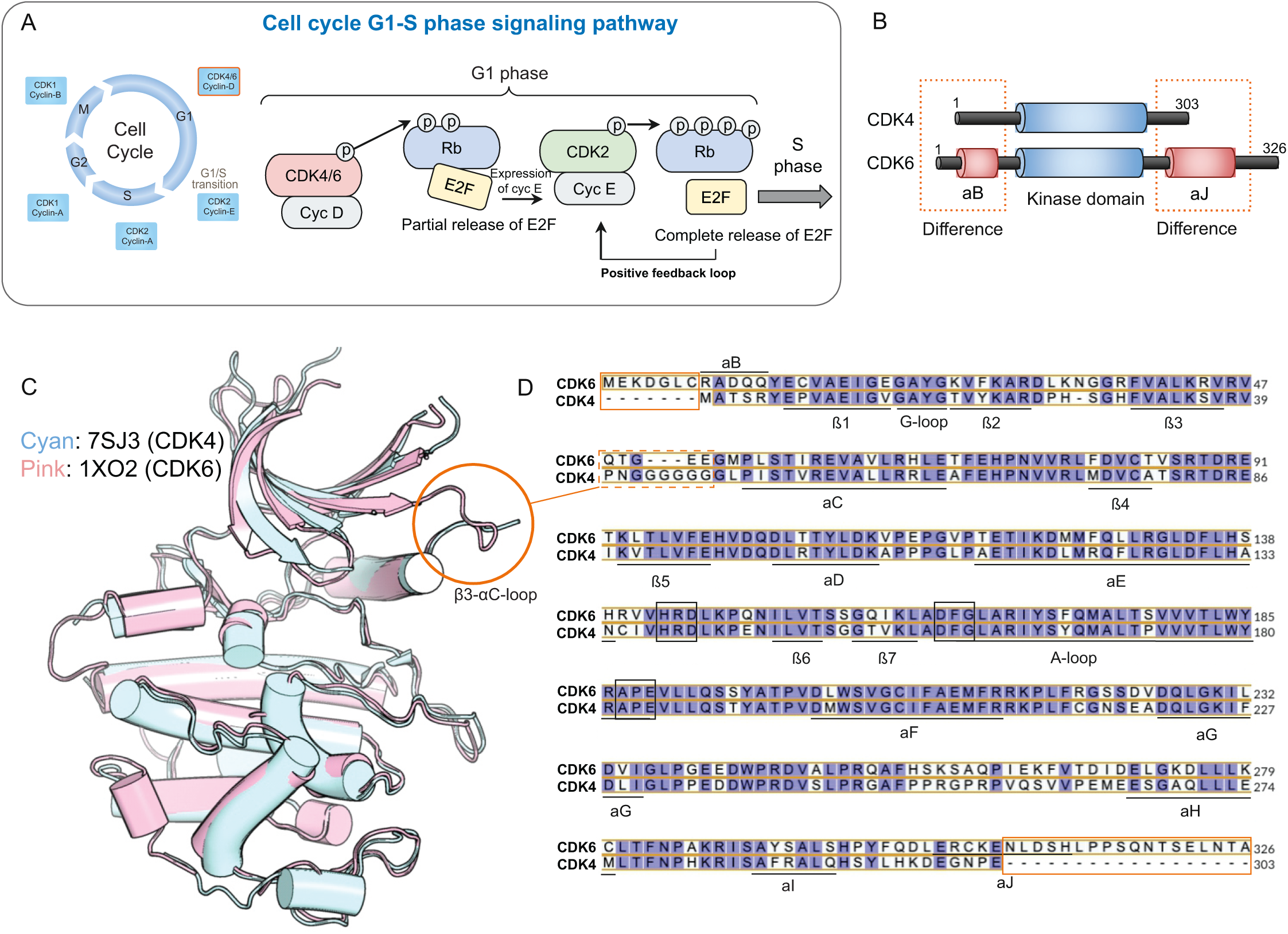
CDK4 and CDK6 domain structures and sequence alignments. (A) Cell cycle and G_1_-S phase signaling pathway. CDK4 and CDK6 are essential in regulating the G_1_ phase of the cell cycle. When complexed with cyclin-D (Cyc D), they phosphorylate Rb, partially releasing E2F transcription factors to induce cyclin-E expression. Cyclin-E/CDK2 then completes Rb phosphorylation, fully releasing E2F, and driving cells into the S phase. The pathway shows the cooperative roles of CDK4/6 and CDK2 in promoting cell cycle progression through a positive feedback loop. (B) Domain structure of CDK4 and CDK6; both CDK4 and CDK6 share a kinase domain but differ in their N-terminal αB-helix and C-terminal αJ-helix regions. (C) Structural alignment of CDK4 and CDK6. The 3D structures of CDK4 (cyan, PDB ID: 7SJ3) and CDK6 (pink, PDB ID: 1XO2) are superimposed, revealing overall similarity with some differences in specific regions. The red circle highlights the β3-αC loop, where structural divergence is observed. (D) Sequence alignment of CDK4 and CDK6 highlights conserved residues (purple background) and divergent residues (white background). Key secondary structures, including α-helices (αB, αC, αD, etc.) and β-strands (β1, β2, etc.), are annotated. Differences in the N-terminal αB helix and C-terminal αJ helix are marked with orange boxes, while variations in the β3-αC loop are emphasized with dashed boxes, corresponding to structural discrepancies shown in panel C. CDK6 contains seven additional residues in the N-terminus and 23 extra residues in the C-terminus, including the αJ helix, which is absent in CDK4.

Previously, our work resolved the question of how cyclin-D/CDK4 evolved to have *slower* catalytic phosphorylation in the long G_1_ phase, whereas cyclin-E/CDK2 evolved to accomplish *fast* phosphorylation of its downstream targets during the short G_1_-to-S transition timescale.[26, 27] We explained the mechanisms, on the conformational level, through which cyclin-D/CDK4 and cyclin-E/CDK2 complexes have adapted to their distinct functional requirements in the G_1_ phase, and G_1_/S phase transition, of the cell cycle. *We clarified why CDK2 can transition from its inactive to the active state faster than CDK4*,[28] *and how the G_1_/S CDK2 complex can maintain faster phosphorylation rate than the G_1_ CDK4 complex*.[29] We suggested the CDK2 structural features which are at play, including (i) stable G-loop, (ii) stable ATP binding site, (iii) stable hydrogen bonds between the hinge region and the adenosine ring of ATP, (iv) stronger protein-protein interactions between the kinase and its activating partner (cyclin) and (v), the stable hydrophobic R-spine that form the backbone of the kinase. These features are necessary but not sufficient for efficient catalysis. Building on these, here we explain how CDK4 and CDK6, despite sharing over 70% sequence identity, similar overall structures, and the same D-type cyclin partner, can exhibit certain distinct functional roles.

Here, we combine biochemical assays and molecular dynamics (MD) simulations analyses to dissect the distinct allosteric regulation of CDK4 and CDK6 and their drug resistance. We attribute the different activities of CDK4 and CDK6 to the local dynamics in the loops (β3-αC, β4-β5, G-loop) in the N-lobe. Our analysis shows that when engaged with cyclin-D, the β3-αC loop allosterically regulates the G-loop, a structure that caps the active site and is essential to kinase activity. In CDK6, the path length between the β3-αC loop and the G-loop is shorter, and the coupling strength between these regions is greater compared to CDK4, enhancing CDK6 activity. Of the few sequence differences between CDK4 and CDK6, the extra sequence in the C-terminus of CDK6 stands out, as it allosterically stabilizes the R-spine, enhancing its catalytic activity. This novel finding extends the long-standing understanding of the roles of the C-terminus in kinase activity,[30, 31] provides a deeper understanding of CDK4/6 function, and may guide the development of next-generation inhibitors.

## 2 Materials and Methods

### Biochemical assays for kinase activity

#### Purification of cyclin/CDK complexes

Human cyclin-D1/CDK4/6 fusion complexes were purified from budding yeast cells (**Table S3**) using a FLAG affinity purification method, modified from a previous protocol.[32] Briefly, N-terminally 3XFLAG-tagged cyclin/CDK fusions with a glycine-serine linker (3XGGGGS) were cloned into high-copy yeast vectors (pRS425) and overexpressed from the GAL1 promoter. The overexpressed 3XFLAG-tagged complexes were then purified by one-step immunoaffinity chromatography using ANTI-FLAG M2 affinity agarose beads (Sigma-Aldrich A2220) and eluted with 0.2 mg/mL 3XFLAG peptide (Sigma-Aldrich F4799).[33]

#### Purification of substrate protein

N-terminally 6His-tagged Rb C-terminal fragment (RbC) recombinant protein (aa: 772-928) was expressed in the *E. coli* strain BL21(DE3)RIL and the purification was performed using cobalt affinity chromatography. Proteins were eluted using buffer containing 25 mM HEPES pH 7.4, 300 mM NaCl, 10% glycerol, and 200 mM imidazole.

#### In vitro kinase assay

*In vitro* kinase assays were performed by incubating 1 µM RbC with cyclin-CDK fusion constructs (in the low nM range) in a reaction mixture containing 100 mM HEPES pH 7.4, 150 mM NaCl, 5 mM MgCl_2_, ∼0.02 mg/ml 3XFLAG peptide, 5% glycerol, 3 mM EGTA, 0.1 mg/ml BSA and 500 μM ATPγS. Reactions were quenched at 8 minutes by addition of EDTA to a final concentration of 25 mM. P-nitrobenzyl mesylate (pNBM) was added to a final concentration of 2 mM and incubated for >1 hour to alkylate thio-phosphorylated proteins. Reactions were separated on 12% SDS-PAGE gels and phosphorylation of substrate proteins was visualized by western blotting. Total protein levels were detected using Revert total protein stain (LiCOR; 926-11010). Western blots were performed using primary rabbit anti-thiophosphate ester antibody (1:2000; Abcam; ab92570) and mouse anti-FLAG antibody (1:5000; Sigma-Aldrich; F1804) coupled with donkey anti-rabbit 800CW and donkey anti-mouse 680CW antibody (1:12000; LiCOR). Fluorescent signal was detected using Odyssey CLX imager (LiCOR) and quantified with the ImageStudio Software (LiCOR). Data were analyzed using GraphPad Prism software (GraphPad). Kinase activity was determined by dividing phosphorylation signal (anti-thiophosphate ester) by both total RbC (detected via total protein staining) and total kinase (anti-FLAG) levels. The activity of all kinases was normalized relative to CDK6^WT^, which was set to 1. Kinase activity, as determined from six independent experiments per enzyme, were compared using one-way ANOVA.

Inhibition assays were conducted as described above, except for the addition DMSO or inhibitor dissolved in DMSO at indicated concentrations. The final DMSO concentration was kept at 10%. Kinase activity was determined by dividing phosphorylation signal (anti-thiophosphate ester) by total RbC (detected via total protein staining) only. Data were fit by non-linear regression using the model “[Inhibitor] vs. response (three parameters)” to determine IC_50_ values. IC_50_ values from three independent experiments per drug-enzyme pair were compared using one-way ANOVA.

### Modeling of CDK complexes and their mutants

We modeled the conformations of the complexes of CDK4 and CDK6 in their cyclin-bound states in the presence of ATP. We constructed the initial coordinates of the active cyclin-D1/CDK4 complex using the crystal structures of cyclin-D3/CDK4 (PDB ID: 7SJ3), replacing the cyclin with cyclin-D1. Cyclin-D1 is modeled using PDB ID 2W9Z, with its C-terminus modeled using homology modeling based on that in cyclin-D3 from 7SJ3. The initial coordinates of cyclin-D1/CDK6 were modeled after the v-cyclin/CDK6 complex (PDB ID: 1XO2) with the cyclin replaced by the same cyclin-D1 as in the CDK4 complex. Both structures are phosphorylated at the active site of the A-loop (pT172 on CDK4, pT177 on CDK6). All missing loops were added using the CHARMM package. We also simulated the active conformations of both CDK4 and CDK6 monomers as well as the unphosphorylated A-loop, from their crystal structures listed above for reference. In all systems, ATP was loaded into both CDKs, replacing the inhibitors in the crystal structures of CDK4 in 7SJ3 and CDK6 in 1XO2 by aligning an ATP-bound crystal structure of CDK2 (PDB code: 1FIN) to both CDKs. The simulated CDK4 and CDK6 systems are listed in **Tables S1, S2**. The solvent was modeled using the TIP3P model with at least 15 Å shell of water in each direction of the periodic boundary box. The charge states of all titratable groups were set to reflect a neutral pH environment, with acidic side chains represented in their negatively charged forms and basic side chains in their positively charged forms. We added Na^+^ and Cl^-^ to neutralize the solvated systems and maintain a physiological salt concentration of 150 mM.

### MD simulation protocols

We performed MD simulations using methods similar to our previous works, in particular utilizing OpenMM.[34–40] The simulation process consisted of several steps up to the production run. We first performed energy minimization using OpenMM’s minimization integrator for 10,000 steps to optimize the systems and remove unfavorable atomic contacts. For each system, we then executed three 2 μs all-atom explicit solvent MD simulations under the NPT ensemble (constant number of atoms, pressure, and temperature) with 3D periodic boundary conditions. We employed OpenMM version 8.1 for the production runs, utilizing the CHARMM 36m all-atom force field. [41–45] Temperature was maintained at 310 K using the Langevin integrator with a friction coefficient of 1 ps^-1^. Pressure was kept constant at 1 atm using a Monte Carlo barostat with an update interval of 25 steps. We applied the SETTLE algorithm to constrain bonds involving hydrogen atoms, allowing for a 2-fs integration time step. Long-range electrostatic interactions were computed using the particle mesh Ewald (PME) method with a grid spacing of 0.1 nm. Short-range non-bonded interactions were calculated using a cutoff of 1.2 nm, with a switching function applied to van der Waals interactions starting at 1.0 nm. The simulations were run on CUDA-enabled GPUs to enhance computational efficiency. Trajectory frames were saved every 100 ps for subsequent analysis. We used MDAnalysis, MDTraj and OpenMM’s built-in analysis tools for post-processing and analyzing the simulation trajectories.[46, 47]

### Principal component analysis of protein dynamics

To analyze the large-scale motions of the protein during the MD simulation, we performed principal component analysis (PCA) using the Bio3D package in R(v4.4.0).[48, 49] PCA was conducted on the Cartesian coordinates of the Cα atoms to capture dominant motions and reduce the dimensionality of the trajectory. A covariance matrix was constructed from the atomic fluctuations of the Cα atoms around their mean positions. The covariance matrix captures the correlated motions of the atoms during the simulation. Next, eigenvalue decomposition of the covariance matrix was performed using the pca.xyz function to obtain eigenvalues and eigenvectors. The eigenvalues represent the variance explained by each principal component (PC), while the eigenvectors describe the direction of the motion in the conformational space. Then, the first few principal components (PCs) were analyzed in detail. Typically, the first few PCs capture the majority of the protein’s collective motion. The contribution of each PC to the total motion was evaluated by the magnitude of the corresponding eigenvalue, and the cumulative variance was plotted to determine how many PCs are needed to describe the essential motions. To further determine the residues that contribute most to the dominant motions, the contribution of individual residues to each PC (loading plot) was calculated using Bio3D. The per-residue contribution was plotted to identify key regions of the protein involved in large-scale conformational changes.

### Community analysis

A network-based community analysis was conducted to identify clusters of residues that move collectively. The dynamic cross-correlation matrix (DCCM) was calculated using Bio3D to quantify correlated and anti-correlated motions between residue pairs. The Cα atom fluctuations from the MD trajectory were used to generate the DCCM. DCCM was then transformed into a distance matrix, and a protein structure network (PSN) was generated where nodes represent residues (Cα atoms), and edges represent significant correlations between them. The edges were weighted based on correlation strength, with a threshold set at 0.5 to filter noise. Communities were detected using the Girvan-Newman algorithm implemented in Bio3D, identifying groups of residues that share similar dynamic properties.

### Binding free energy calculations

To quantitatively evaluate the binding affinities and specificities between cyclin-D/CDK4 and cyclin-D/CDK6 complexes, we determined the binding free energies using the molecular mechanics/generalized Born surface area (MM/GBSA) approach, implemented in AmberTools and Amber.[50] This method allows us to assess the interactions between cyclin-D and CDK4 or CDK6 by calculating the binding free energies from MD simulations of the complexes. The total binding free energy (ΔG_b_) is calculated as the sum of the molecular mechanics gas-phase energy (ΔE_MM_), solvation free energy (ΔG_sol_), and entropy contributions (-*T*ΔS), where *T* is the temperature (in Kelvin):

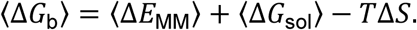

The ensemble averages, denoted by the angle brackets, were calculated over the final 1 μs of equilibrated configurations from the MD simulations. The components of the gas-phase molecular mechanics energy (ΔE_MM_) include the internal energy (ΔE_inter_), electrostatic interactions (ΔE_elec_), and van der Waals interactions (ΔE_vdW_):

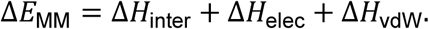

The solvation free energy (ΔG_sol_) consists of the polar and nonpolar contributions, computed using the generalized Born (GB) model in AmberTools with the igb=5 parameter (GBneck2 model) for the polar solvation energy and a surface area-based model for the nonpolar component. The nonpolar contribution is calculated using the linear combination of pairwise overlap (LCPO) model:

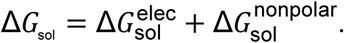

The entropy contributions (-*T*ΔS) to the binding free energy were estimated using normal mode analysis (NMODE) performed within AmberTools. These include translational (*T*ΔS_trans_), rotational (*T*ΔS_rot_), and vibrational (*T*ΔS_vib_) components. The vibrational contributions were computed from quasi-harmonic approximations based on the trajectory snapshots:

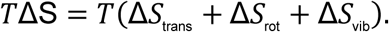

The net binding free energy (ΔG_b_) was determined by calculating the difference between the binding free energy of the bound complex and the sum of the free energies of the unbound CDK and cyclin proteins:

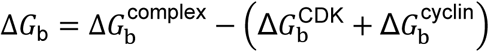

### Allosteric pathway analysis

To investigate allosteric pathways in our protein systems, we conducted a dynamical network analysis using the Bio3D package,[48] implementing the weighted implementation of suboptimal paths (WISP).[51] Each amino acid residue was treated as a node, with edges connecting residues whose Cα atoms were within a 5 Å threshold. Edge weights, -ln(*c_ij_*), where *c_ij_* is the correlation between residues forming the edge, are inversely proportional to the dynamic correlations calculated from MD simulations, indicating residue coupling strength. We selected source and sink residues based on dynamic cross correlation and their functional relevance, typically involving key binding regions undergoing conformational changes during allosteric modulation. The path length is defined as the sum of edge weights along a path, with longer paths indicating weaker coupling between source and sink residues.

Using WISP, we calculated the top 100 suboptimal paths between source-sink pairs, capturing multiple communication routes and potential alternative pathways for signal propagation. Path lengths and their distributions were analyzed to assess coupling strength across the network. These analyses inform the protein’s allosteric landscape, showing key residues mediating connectivity between functionally important sites.

### Quantification and statistical analysis

For statistical robustness, each complex was subjected to three independent MD simulations, with the final binding free energy values representing the average over these runs. The convergence of the simulations was assessed by monitoring the potential energy, electrostatic energy, and the root-mean-square deviation (RMSD) of the Cα atoms, ensuring that equilibrium was reached. To improve statistical accuracy, only the last 1 μs of each trajectory was used for energy calculations. Result analyses were carried out using OpenMM,[44, 45] VMD,[52] PyMOL,[53] MDAnalysis,[46] and custom Python scripts. All free energy results were reported as mean ± standard deviation across the independent simulation replicates.

## 3 Results

### CDK6’s longer C-terminus and CDK4’s elongated β3-αC loop define their structural contrasts

A high degree of functional and structural similarity has resulted in CDK4 and CDK6 often being referred to as “CDK4/6”. However, both possess unique evolutionarily conserved structural features **(Fig. S2).** CDK6 has extended N- and C-termini, each containing an additional α-helix (αB and αJ) (**Fig. 1B-D)**. While the function of these extensions has yet to be investigated, in transcriptional CDKs, C-terminal extensions help to shape the active site.[54] CDK4 has a longer loop connecting its β3-strand to its αC-helix (β3-αC loop), ^40^PNGGGGGG^47^ compared to ^48^QTGEE^52^ in CDK6. Salt bridge formation between these two structural elements is required to form the ATP binding pocket, thus their relative positions are of critical importance in kinase activation.[28, 55] For example, decreasing the length of the β3-αC loop has been shown to lock BRAF in an active conformation and reduce inhibitor efficacy.[56] By contributing to distinct allosteric networks, these subtle structural variations may impart the distinct regulatory and functional properties of CDK4 and CDK6 in cell cycle control. This work explores the significance of these allosteric structural differences in relation to their activity and functional implications.

### Enhanced flexibility in CDK4 loops and C-lobe correlates with reduced activity

Despite only minor differences in sequence and structure between CDK4 and CDK6, biochemical assays reveal that wild-type CDK6 exhibits more than 10-fold greater kinase activity toward the Rb C-terminus (RbC) when compared to wild-type CDK4 (**Fig. 2A and S3**). To understand the structural origins of this difference, we performed principal component analysis (PCA) on the MD trajectories of cyclin-D/CDK4 and cyclin-D/CDK6. We denote these two systems CDK4^WT^ and CDK6^WT^. Analysis shows vastly different conformational clustering patterns for each kinase, with conformational ensembles depicted by PCA revealing distinct local fluctuation patterns for CDK4^WT^ and CDK6^WT^ in the first principal component in particular (**Fig. 2B**). The first two principal components explain a larger portion of the variance in CDK4^WT^ (39%) than in CDK6^WT^ (34%), suggesting greater conformational flexibility in CDK4^WT^ (**Fig. 2C**). Residues in the β3-αC and β4-β5 loops display high degrees of conformational variability, as indicated by their contribution to PC1 (**Fig. 2D**). Conformational ensembles play a crucial role in determining function. [57, 58] Despite their sequence homology, CDK4 and CDK6 occupy different conformational spaces, likely reflecting their non-redundant functional roles in cell cycle progression.[16, 54, 59] This is consistent with findings that CDK6 has a more stable αC-helix conformation compared to CDK4, potentially explaining its higher basal activity.[60] The dynamic cross-correlation map for all residue pairs shows distinct patterns of intramolecular communication within CDK4^WT^ and CDK6^WT^ (**Fig. 2E**). CDK4^WT^ shows more extensive anti-correlations between the N- and C-lobes. Excessive anti-correlation indicates less stable structure leading to lower efficiency compared to other cell cycle CDKs.[29]

**Figure 2.**
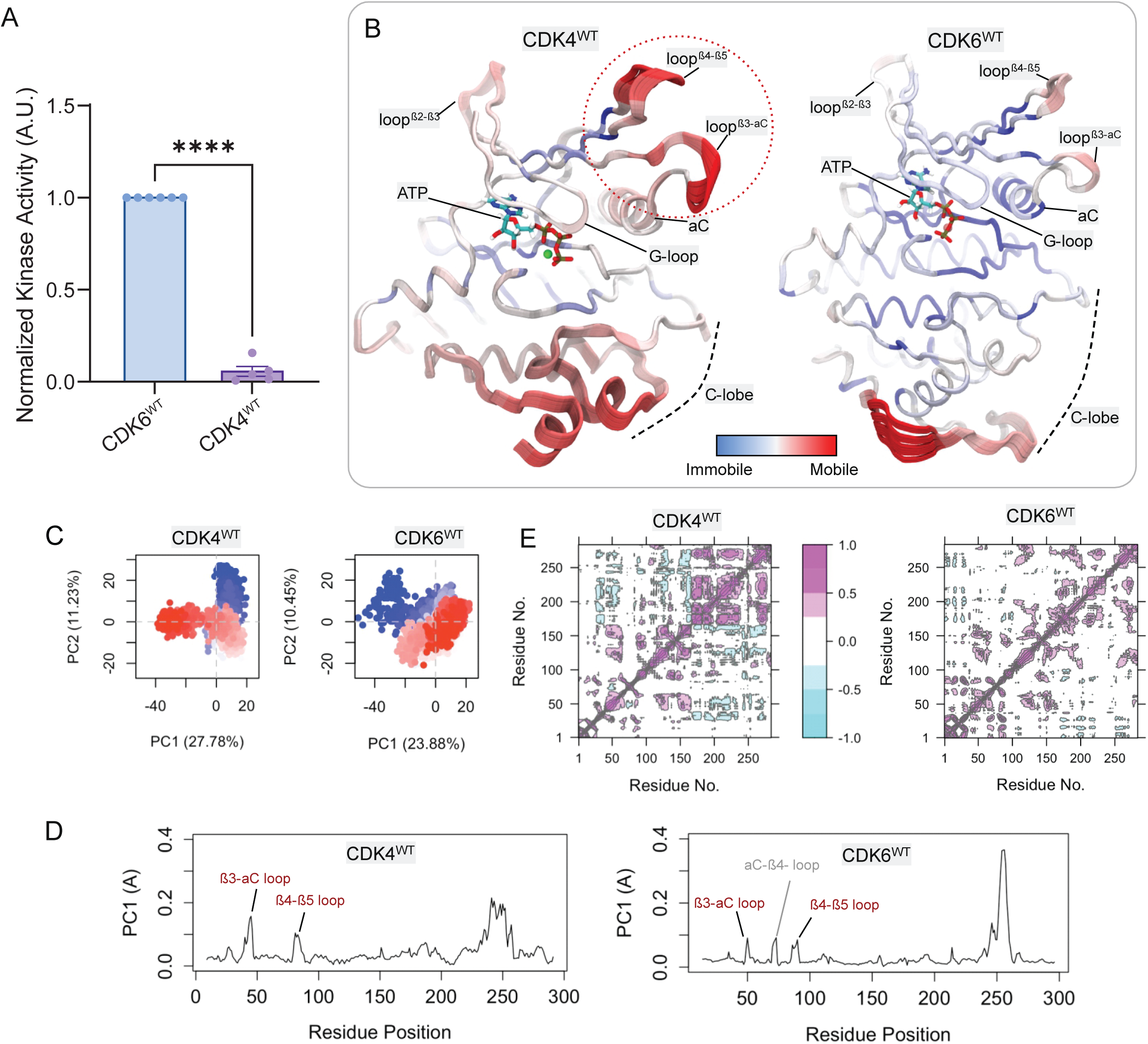
CDK6^WT^ is more active than CDK4^WT^, in part due to its lower local dynamics in β3-αC and β4-β5, and G-loops. (A) Normalized kinase activity of cyclin D1-CDK6^WT^ and cyclin D1-CDK4^WT^ fusions toward RbC. (N = 5-6; ****P<0.0001) (B) Tube representations of the first principal component (PC1) from PCA of CDK4^WT^ and CDK6^WT^. Color gradients represent mode mobility, from immobile (blue) to highly mobile (red). Notable dynamic regions include the β3-αC, β4-β5, and β2-β3, and G-loops. ATP is shown bound in the active site. The binding cyclins are not displayed for visual clarity. (C) PCA plot for the first and second principal component (PC1 and PC2) of CDK4^WT^ and CDK6^WT^, show distinct clustering patterns. For CDK4^WT^, PC1 and PC2 account for 27.78% and 11.23% of the total variance, respectively. For CDK6^WT^, PC1 and PC2 explain 23.88% and 10.45% of the total variance, respectively. Each dot represents one snapshot from the trajectories in the plot. The color ranging from blue to red represents the trajectory frame from beginning to end. (D) PC1 loading plots for CDK4^WT^ and CDK6^WT^. The fluctuation profiles identify regions of high conformational variability, particularly in the β3-αC and β4-β5 loops. β3-αC loop has more significant contribution to the PC1 than that of CDK6. (E) Dynamic cross-correlation maps for CDK4^WT^ and CDK6^WT^. Positive correlations (magenta) and anti-correlations (cyan) highlight coupled motions between specific regions.

### Longer β3-αC loop in CDK4 disrupts coordinated kinase dynamics and activity

CDK4^WT^ has a longer and more flexible loop connecting the β3-strand to the αC-helix relative to CDK6^WT^ (**Fig. 3A and 3B** and **Fig. S4**). During the transition from the inactive to the active state, the longer loops in CDK4 block the inward movement of the αC-helix upon cyclin binding, contributing to its slower activation compared to CDK2.[28] However, once CDK4 crosses the activation barrier and becomes the active complex (as in the state studied here), the β3-αC loop also plays a role in modulating the dynamics of the complex by decoupling the correlated motion of the αC-helix and the G-loop, hence decreasing the activity. The binding free energy indicates a stronger affinity between cyclin-D and CDK6^WT^ compared to CDK4^WT^ (**Fig. 3C**) due to more extensive contacts with cyclin-D (**Fig. S5**), resulting in the stronger coupling. This stronger association may contribute to the higher catalytic efficiency of CDK6, analogous to highly efficient cyclin-E and CDK2 having a greater binding affinity than cyclin-D and CDK4 in our previous study.[29] The flexible loop decreases binding affinity since flexibility increases the entropic contribution to the binding free energy, which decreases the binding affinity (**Fig. S6**).[61]

**Figure 3.**
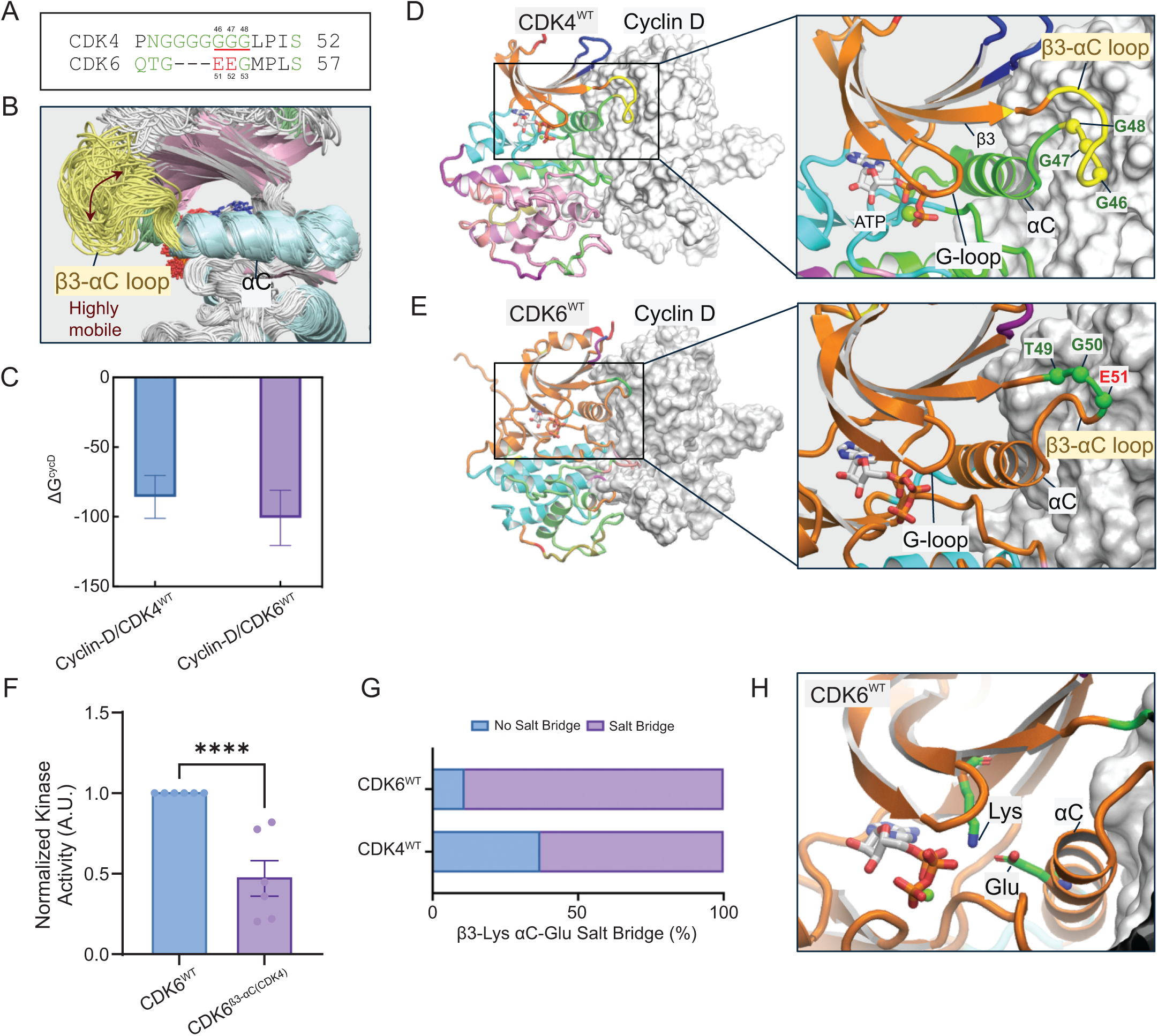
Longer β3-αC loop of CDK4 is the key in the binding interface between CDK4 and cyclin-D and decouples N-lobe β-strands and αC-helix, negatively regulating their activities. (A) Sequence alignment of the β3-αC loop region in CDK4 and CDK6. This panel shows the β3-αC loop of CDK4 has a significantly longer sequence of glycine residues than CDK6. (B) Structural overlay of flexible β3-αC loop region from the trajectories of CDK4^WT^. The overlay demonstrates the highly mobile nature of the β3-αC loop in CDK4^WT^. (C) Binding free energy (ΔG^CycD^) of cyclin-D with CDK4^WT^ and CDK6^WT^. The bars represent the calculated ΔG for the binding of cyclin-D to wild-type CDK4 (left, blue) and wild-type CDK6 (right, purple). More negative values indicate stronger binding, with error bars representing the standard deviations. Community network of (D) cyclin-D/CDK4^WT^ and (E) cyclin-D/CDK6^WT^ complexes. In the CDK4 complex (D), the G-loop and αC-helix are in different communities, whereas in the CDK6 complex (E), they are in the same community. Different colors of the cartoon representation of CDKs represent distinct communities. (F) Normalized kinase activity of cyclin D1-CDK6^WT^ and cyclin D1-CDK6^β3-αC(CDK4)^ fusions toward RbC. (N = 6; ****P<0.0001) (G) Probability of β3-Lys and αC-Glu salt bridge formation. This bar graph shows the higher probability of the formation of the β3-Lys and αC-Glu salt bridge in CDK6^WT^ compared to CDK4^WT^. The higher probability in CDK6^WT^ suggests more stable active conformation. (H) Snapshot highlighting the Lys-Glu salt bridge in CDK6^WT^. This conformation is characteristic of the active conformation of kinases.

We next used community analysis to understand how the longer β3-αC loop in CDK4 affects the coordinated rearrangement of structural components essential for kinase activation. Community analysis, which segments proteins into smaller, interconnected groups of residues that move together as semi-rigid bodies, offers valuable insights into the cooperative dynamics of protein regions and the propagation of conformational changes throughout the structure. [14, 62–64] Structural analysis of the cyclin-D/CDK6^WT^ complex places the G-loop and αC-helix in the same structural community, indicating coordinated movement between both structures (**Fig. 3E**). These structural elements play critical roles in facilitating catalysis, with the G-loop capping the active site and αC-helix ensuring that the kinase remains in an active state through its interactions with the β3-strand. Coordinated motion between the G-loop and αC-helix is associated with enhanced kinase activity, as seen in eukaryotic protein kinases.[65] Cyclin-D/CDK4^WT^ lacks this coordination, as community analysis splits the G-loop and the αC-helix into different structural communities (**Fig. 3D**). This is at least partially due to the longer β3-αC loop in CDK4, as swapping the β3-αC loop in CDK6 for that of CDK4 (CDK6^β3-αC(CDK4)^) reduces catalytic activity by 50% and results in communities like those observed for CDK4^WT^ (**Fig. 3F and S7**). Thus, this difference in dynamic organization may explain the higher basal activity observed in CDK6.

Finally, we assessed how the structural differences between CDK4 and CDK6 affects β3-strand αC-helix contacts. In active cyclin-CDK complexes, the β3-strand lysine (K35 in CDK4, K43 in CDK6) forms a salt bridge with the αC-helix aspartate (E56 in CDK4, E61 in CDK6) (**Fig. 3G**), creating a link to the G-loop, thereby stabilizing both elements. Our simulations reveal that CDK6^WT^ has a far higher probability of salt bridge formation than CDK4^WT^, suggesting a more efficient catalysis-ready conformation. Mutations affecting this residue disrupt the relative positioning of the G-loop and αC-helix, altering the active site and reducing catalytic efficiency.[65, 66] Given the importance of active site precision for efficient catalysis[66], coordinated motion between the G-loop and the αC-helix likely optimizes this conformation (**Fig. 3H**), leading to enhanced kinase activity.

### Stronger allosteric coupling in CDK6 stabilizes active site for efficient catalysis

We next used network analysis to define the mechanism of allosteric regulation between the β3-αC loop and the G-loop. This network-based approach allows us to understand how signals travel through the kinase and how different structural elements work together *dynamically*. The strength of the connection between two residues is based on how correlated their motions are: if they tend to move together, they form a path and are strongly connected. We focused on the allosteric signaling pathway linking the β3-αC loop (source) to the G-loop (sink) with source and sink residues selected based on the dynamic cross-correlation map (**Fig. 2E**). The selected source residues are G46 of CDK4 and E51 of CDK6 in the β3-αC loop and the sink residues are A16 of CDK4 and A23 of CDK6 in the G-loop.

The allosteric signaling path in CDK4^WT^ is long and meandering, starting from its extended β3-αC loop and progressing through the β3-strand, β2-strand, and β1-strand before reaching to the G-loop (**Fig. 4A and C**). In contrast, the CDK6^WT^ allosteric pathway is more direct, with the allosteric signal only passing through a small portion of the β3- and β2-strands without influencing the β1-strand (**Fig. 4B and E**). To investigate how strongly the residues along these paths are connected, we calculated the length of the allosteric path. The path length is the sum of edge weights in a network path connecting two sites, where each edge weight is typically calculated as -ln(|*c*_ij_|) with *c*_ij_ being the correlation between connected residues *i* and *j*, rather than the physical distance between the source and sink residues, or the number of residues between the source and sink along a path. The path length distribution in CDK4^WT^ (peak at ∼3.44) indicates slightly longer paths with weaker coupling between the source and sink residues compared to CDK6^WT^ (peak at ∼3.1) (**Fig. 4D and F**). The shorter allosteric path length in CDK6^WT^ indicates stronger correlated motions between the β3-αC loop and the G-loop, potentially positively modulating the kinase activity. The different allosteric pathways can be attributed to the different length of the β3-αC loop. CDK4^WT^ has a longer β3-αC loop than that of CDK6^WT^ (**Fig. 1D)**. Loop swapping of CDK6^β3-αC(CDK4)^ by adopting the β3-αC loop of CDK4 results in longer allosteric paths, resembling those observed for CDK4^WT^ (**Fig. S7**).

**Figure 4.**
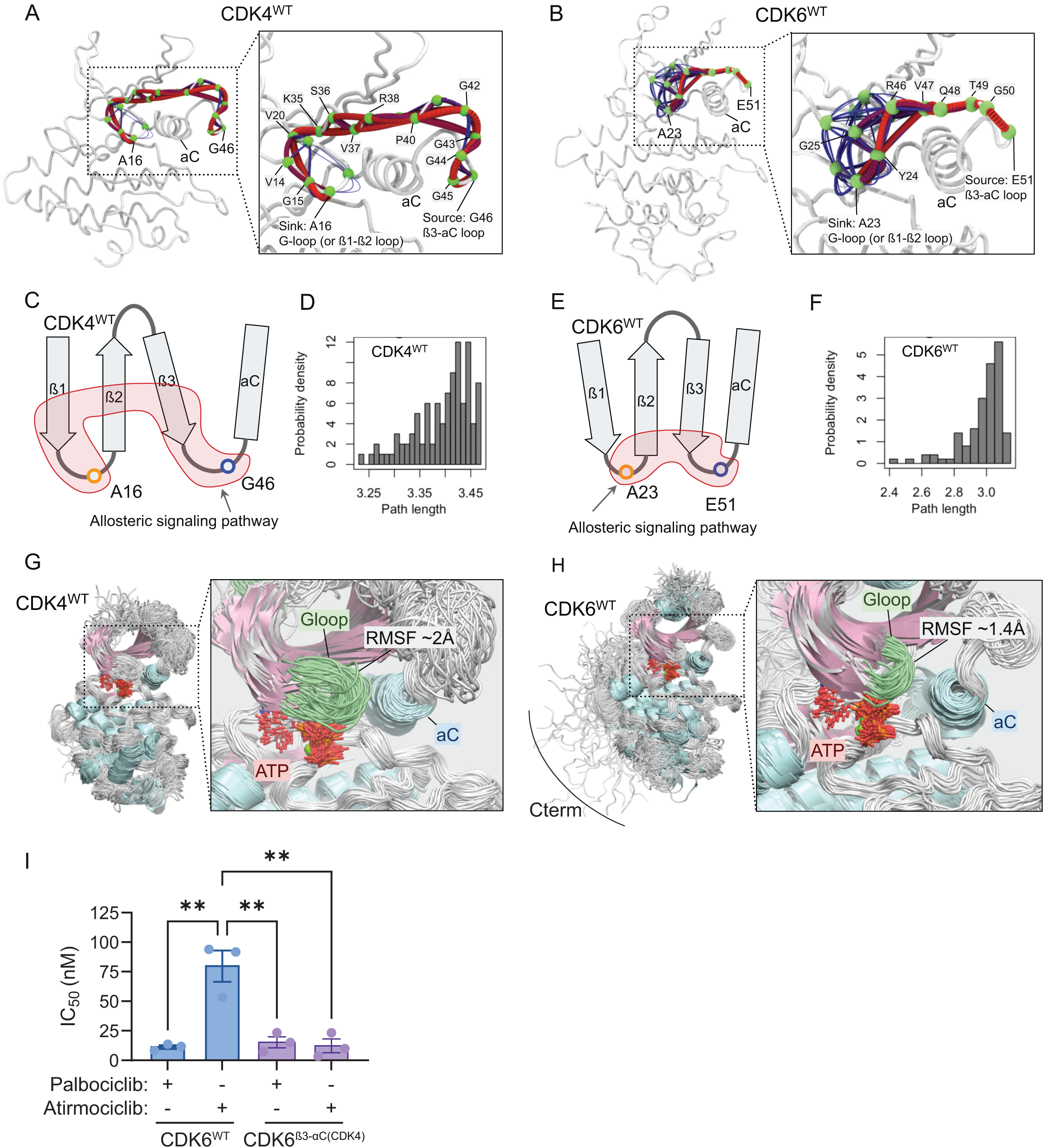
Allosteric signaling pathways between the β3-αC loop and the G-loop. Optimal (red lines) and sub-optimal (blue lines) allosteric paths for (A) CDK4^WT^, and (B) CDK6^WT^. The allosteric signaling pathway starting from the β3-αC loop to G-loop of CDK4 follows a longer path than that of CDK6. The green spheres are α-carbon of residues and represent nodes in the allosteric pathways. (C) Schematic illustration of the optimal allosteric paths and (D) path length distribution between the source and sink residues for CDK4^WT^. (E) Schematic illustration of the optimal allosteric paths and (F) path length distribution between the source and sink residues for CDK6^WT^. Shorter path length suggests a stronger allosteric pathway between the source and sink. Overlay of simulation trajectories for (G) CDK4^WT^ and (H) CDK6^WT^. G-loop (green) of CDK4 is longer than that of CDK6 and is more dynamic. (I) IC_50_ values for palbociclib and atirmociclib mediated inhibition of cyclin D1-CDK6^WT^ (11.17 ± 1.51 nM and 79.69 ± 13.23 nM) and cyclin D1-CDK6^β3-αC(CDK4)^ (15.22 ± 4.66 nM and 12.18 ± 5.84 nM). (N = 3; **P<0.005)

The G-loop connecting the β1- and β2-strands contains a conserved GxGxxG motif and is located on top of ATP, which is important for ATP positioning. Superimposition of the conformations from the simulated trajectories shows that the G-loop of CDK4^WT^ has greater flexibility (**Fig. 4G**) compared to that of CDK6^WT^ (**Fig. 4H**), as indicated by the residual RMSFs (**Fig. S9**). These observations could provide an explanation for the different responses of CDK4 and CDK6 to certain inhibitors, such as the CDK4-selective inhibitor atirmociclib (PF-07220060), which exhibits a remarkably high selectivity for CDK4 over CDK6. The enhanced flexibility of the CDK4 G-loop in part allows for more efficient binding of the bulky isopropyl alcohol [(CH_3_)_2_COH] group in this inhibitor (**Fig. S10**).[67] Indeed, replacing the β3-αC loop in CDK6 with that of CDK4 (CDK6^β3-αC(CDK4)^) significantly increases its sensitivity to this CDK4-selective inhibitor (**Fig. 4I and S11)**.

### CDK6 C-terminus stabilizes the R-spine, enhancing catalytic activity

Relative to CDK4, CDK6 has an extended C-terminal tail, a structural feature known to regulate the kinase activity. For example, the C-terminal tail of PKA supports its active conformation.[30, 31] Some CDKs are known to have flexible C-terminal tail that helps to stabilize the N-lobe by folding onto it, enclosing the ATP in its active site.[54] However, it is not known how the C-terminal tail of CDK6 affects its conformational dynamics. To elucidate the role of the C-terminus, we simulated CDK6 with a truncated C-terminal tail (CDK6^ΔCterm^) and calculated the cross-correlation of atomic fluctuations. The dynamic cross-correlation map reveals highly correlated residue motions between the N- and C-lobes of CDK6^WT^ (**Fig. 5A and C**). In contrast, these residue motions are uncorrelated in CDK6^ΔCterm^ (**Fig. 5B**). Instead, correlation is primarily observed within the C-lobe (**Fig. 5D**). We hypothesize that in CDK6^ΔCterm^, the absence of the correlated residue pairs between the N- and C-lobes renders the inter-lobe motions rigid, hindering the opening and closing of the active site and resulting in lower activity. Consistent with this, biochemical assays with CDK6^ΔCterm^ show reduced kinase activity compared to CDK6^WT^, resulting in a lower rate of RbC phosphorylation (**Fig. 5E**). One explanation for the reduced activity of CDK6^ΔCterm^ is that the C-terminal deletion disrupts the coordinated movements of the regulatory-spine (R-spine) of CDK6 (**Fig. 5F**). Once the R-spine is disrupted, the kinase is less likely to maintain its stability and resulting in reduced activity.[68] In CDK6^WT^, we observed multiple strong and short allosteric signaling paths connecting the C-terminal edge of the kinase domain to an R-spine residue (**Fig. 5G**), but this is not the case in CDK6^ΔCterm^ (**Fig. 5H**). CDK6^ΔCterm^ disrupts the intrinsic dynamics of the kinase, resulting in longer allosteric paths from the C-terminus through both αH- and αF-helices to the R-spine. This is in part due to the high mobility of the truncated C-terminal tail (**Fig. S12**), which increases the dynamics of the C-lobe, breaking the intrinsic coordinated motions within the kinase that separate R-spine into three distinct communities (**Fig. 5F**). Our finding represents a novel mechanism of kinase activity in which the C-terminal tail plays a direct role in modulating the opening and closing of the C- and N-lobes through R-spine dynamics.

**Figure 5.**
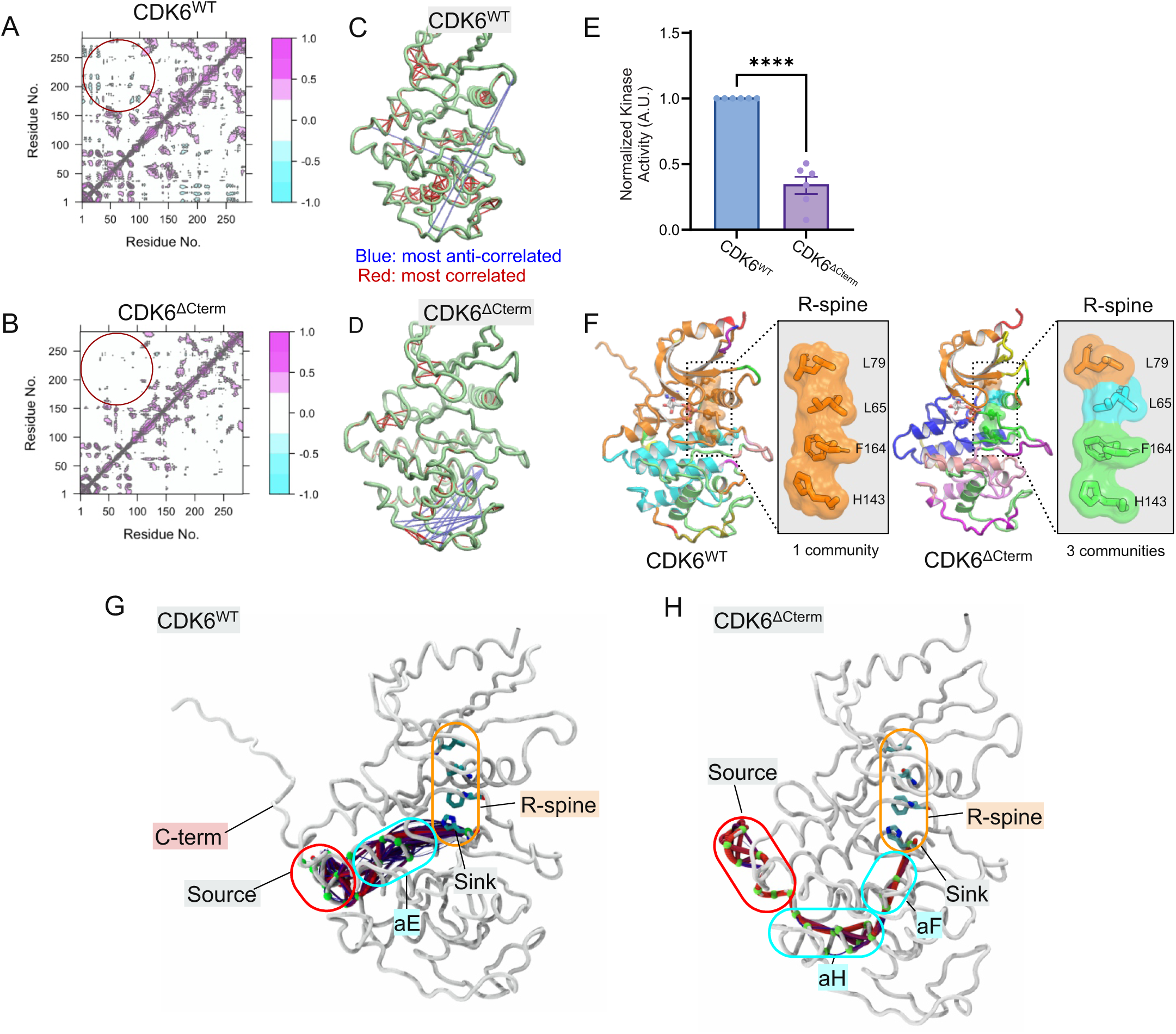
C-terminus modulates the C- and N-lobe opening and closing of CDK6 facilitating more efficient catalysis. Dynamic cross-correlation maps of residue for (A) CDK6^WT^ and (B) CDK6^ΔCterm^. The maps use a color scale from cyan (-1.0, anti-correlated) to magenta (+1.0, correlated) to show how different protein residues move in relation to each other. A key difference is highlighted in the circled region, showing distinct correlation patterns between N-lobe (residues 1-100) and C-lobe (residues >100) movements in the two kinases. Tube representations of the most correlated and anti-correlated motion in (C) CDK6^WT^ and (D) CDK6^ΔCterm^. These panels show the regions of CDK6^WT^ and CDK6^ΔCterm^ with the most correlated (positive correlations, red) and anti-correlated (negative correlations, blue) motions, showing how the truncation of the C-terminus alters these dynamics. (E) Normalized kinase activity of cyclin D1-CDK6^WT^ and cyclin D1-CDK6^ΔCterm^ fusions toward RbC. (N = 6; ****P<0.0001) (F) Structural community analysis of CDK6^WT^ and CDK6^ΔCterm^ focusing on the regulatory spine (R-spine). In CDK6^WT^, the R-spine residues (L79, L65, F164, H143) form a single cohesive community (orange), while in CDK6^ΔCterm^, these residues are dispersed across three different communities (colored differently), suggesting compromised structural integrity. Allosteric pathway analysis in (G) CDK6^WT^ and (H) CDK6^ΔCterm^. The graphs trace the communication pathway (highlighted in various colors) from the source residue E308 in the C-terminal region to the sink residue H143 in the R-spine. CDK6^WT^ shows a direct pathway through the αE-helix, while CDK6^ΔCterm^ exhibits a longer route through multiple helices (αH and αF), indicating altered internal dynamics due to C-terminal truncation.

## 4 Discussion

Though largely identical, CDK4 and CDK6 evolved distinct activities in regulating the G_1_ progression. This distinction is particularly relevant to CDK4/6 inhibitor-resistant cancer cells, as well as in adult tissues and HSCs. The key question is how the overexpressed CDK6 in cancer cells enables cell cycle progression in lieu of CDK4, and what molecular mechanisms differentiate the higher activity of CDK6 from that of CDK4. Explaining these differences at the conformational level could clarify the shifts towards the CDK6 complex for cell cycle progression in resistant cancer cells, thereby bypassing CDK4/6 inhibition.

Our experimental results indicate that CDK6^WT^ is more active than CDK4^WT^ and CDK6^ΔCterm^. The latter, which lacks the C-terminal part of CDK6 is more similar to CDK4 in terms of activity. We use MD simulations to explain the molecular origin of their catalytic differences. The higher activity of CDK6^WT^ can be attributed to several factors, including the allosteric role of the β3-αC loop upon cyclin binding. A longer β3-αC loop in CDK4 results in a *disjointed* kinase, while a shorter loop in CDK6 is better at facilitating *coordinated* motion for efficient catalysis. Furthermore, the weaker coupling between the G-loop and the β3-αC loop in CDK4^WT^ allows decoupling of the β-stands (β1-β5) and the αC-helix. This results in insufficient stabilization of ATP and, consequently, lower activity due to the longer β3-αC loop of CDK4^WT^. In contrast, CDK6^WT^ has a shorter β3-αC loop, resulting in stronger coupling between the G-loop and the β3-αC loop. This enhanced coupling leads to better stabilization of ATP and higher catalytic activity. Secondly, the unstructured C-terminus of CDK6^WT^ acts on the kinase allosterically, stabilizing the R-spine for more efficient catalysis. In contrast, CDK4^WT^ lacks this C-terminus, leading to a less stable R-spine, as shown in our previous works.[29] Thirdly, the dynamics of CDK4^WT^ reveal a significant degree of opening and closing of the N- and C-lobes, leading to an unstable structure as evidenced by the unstable R-spine.[29] This instability likely contributes to the slower catalytic activity observed for CDK4, as the excessive conformational changes typically hinder efficient substrate binding and phosphoryl transfer (**Fig. 6**). In contrast, as discussed in our previous work,[29] CDK2^WT^ displayed a relatively stable structure with less dynamic opening and closing motions of the N- and C-lobes for ATP binding, resulting in higher catalytic activity. We suspect that CDK6^WT^ may exhibit the right balance of N- and C-lobe movements between CDK2 and CDK4, and a coordinated and efficient catalytic process. The ability of CDK6^WT^ to maintain a dynamic yet stable conformation is likely responsible for its higher activity compared to both CDK4^WT^ and CDK6^ΔCterm^.

**Figure 6.**
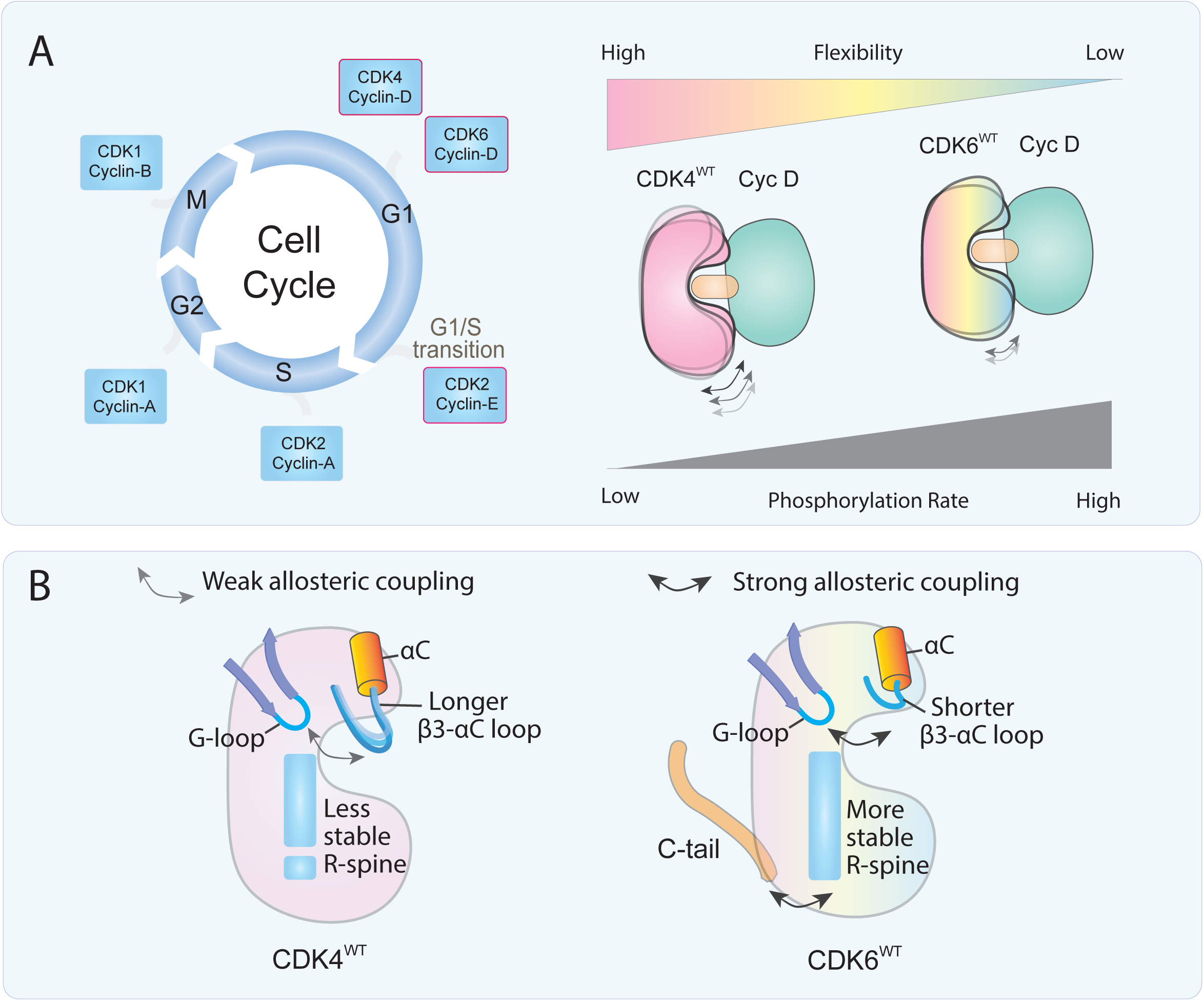
CDK4 and CDK6 dynamic divergence gives rise to their distinct activity. (A) Schematic of cell cycle showing CDK-cyclin complex activity at different phases (G_1_, S, G2, M). Gradient scales show that CDK4^WT^ exhibits higher flexibility but lower phosphorylation rate, while CDK6^WT^ shows lower flexibility but higher phosphorylation rate. This inverse relationship between flexibility and catalytic efficiency reflects the evolutionary adaptation of CDK6 for rapid cell division in stem-like cells like ST-HSC cells. (B) Structural comparison of CDK4^WT^ and CDK6^WT^ highlighting key differences in their regulatory mechanisms. CDK4^WT^ has a longer β3-αC loop that results in weak allosteric coupling between αC-helix and the G-loop, while CDK6 has a shorter β3-αC loop with a stronger allosteric coupling, and its C-terminus stabilizes the R-spine for more efficient phosphorylation.

These findings suggest that achieving optimal kinase function requires the right balance of conformational plasticity and show that the evolution of CDK6, as exemplified by its role in fast-proliferating short-term HSCs, has favored allosteric networks that enhance its activity relative to CDK4. In CDK4/6i-resistant cancer cells, CDK6 is often overexpressed. When complexed with CIP/KIP family inhibitors, such as p18 or p27, CDK6 becomes insensitive to the inhibitors, in part due to the allosteric mechanisms it has acquired through evolution.[22, 60] These mechanisms include the longer C-terminus, which stabilizes the R-spine, and stronger coupling between the β3-αC loop and the αC-helix upon cyclin binding.

The divergence in sequence, function, and activity of CDK4 and CDK6 is not unique in evolution, and our results show that such differences provide opportunities for designing inhibitors that selectively target one kinase but not the other. Notably, FGFR1 and FGFR2 follow a similar pattern.[69, 70] Despite their high sequence similarity, these kinase pairs exhibit different catalytic efficiencies and regulatory mechanisms. The different dynamics of FGFR1 and FGFR2 have been leveraged in the design of the inhibitor lirafugratinib to selectively target FGFR2.[70] The G-loop (or P-loop) in FGFR1 is highly flexible, rapidly transitioning between an extended conformation and a more contracted state. In contrast, the G-loop in FGFR2 maintains a relatively extended conformation, which is far less dynamic than that of FGFR1, allowing efficient covalent binding of lirafugratinib to C491 in FGFR2. Although lirafugratinib stabilizes the G-loop conformation in FGFR1, the covalent bond is too far away from C488 for efficient covalent binding. These examples show the importance of considering protein dynamics in drug design.

Closely related kinases, such as CDK4 and CDK6, often have different allosteric mechanisms that can potentially be exploited to selectively target one over the other. For example, AKT1 is more biochemically active and responsible for the tumor initiation, while AKT2 is mainly responsible for tumor progression and metastasis.[71] Their divergence is in part allosterically regulated by their differences in the PH domain.[72]

Here, we discover two novel allosteric mechanisms that CDK4 and CDK6 utilize to differentially regulate their activity: allosteric signaling between the β3-αC loop and the G-loop, “sensing” cyclin binding, and the C-terminus of CDK6, modulating its activity through coordinated R-spine movement. The weak allosteric coupling of β3-αC loop and G-loop in the CDK4 offers a more flexible G-loop compared to CDK6. These allosteric mechanisms may also in part explain why overexpression of CDK6, rather than CDK4, leads to drug resistance. This is because CDK6 may be more efficient at promoting cell cycle progression even when partially inhibited, and the relatively inflexible G-loop of CDK6 is less likely to allow binding of bigger ligands such as inhibitors, while simultaneously allowing binding and unbinding of ATP, facilitating ATP hydrolysis.[22, 73] To overcome CDK6 overexpression in drug resistance to CDK4/6i, proteolysis-targeting chimeras (PROTACs) and molecular glue degraders have been proposed.[59, 73] This strategy is highly promising as it potentially eliminates the overexpressed CDK6 through its targeted degradation, rather than merely inhibiting its activity.

## 5 Conclusions

Our combined approach, leveraging molecular simulations and biochemical assays, uncovers new molecular mechanisms underlying the allosteric differences between the closely related CDK4 and CDK6.Two allosteric networks are identified: one in the N-lobe (connecting the β3-αC loop and G-loop) and another in the C-lobe (linking the C-terminus and R-spine). In CDK6 these networks have shorter paths or stronger coupling than those in CDK4, providing favorable conditions for higher kinase activity. These findings reveal how these kinases evolved to fulfill distinct functional roles in different cell types and offer new mechanistic details into their allosteric regulation and inhibitor specificity.

## Acknowledgments

We thank members of the Kõivomägi and Nussinov lab for the discussions and comments on this work. This project was supported in whole or in part by federal funds from the National Cancer Institute, National Institutes of Health, under contract HHSN261201500003I and ZIABC012133. The contents of this publication do not necessarily reflect the views or policies of the Department of Health and Human Services, nor does mention of trade names, commercial products, or organizations imply endorsement by the U.S. government. This research was supported [in part] by the Intramural Research Program of the NIH, National Cancer Institute, Center for Cancer Research. All simulations had been performed using the high-performance computational facilities of the Biowulf cluster at the National Institutes of Health, Bethesda, MD (https://hpc.nih.gov/).

## Author contributions

**Wengang Zhang**: Conceptualization, Methodology, Software, Visualization, Investigation, Data collection and curation, Writing – original draft & editing **Devin Bradburn**: Conceptualization, Methodology, Investigation, Data collection and curation, Visualization, Writing – original draft & editing **Gretchen Heidebrink**: Investigation **Yonglan Liu** Methodology, Software **Hyunbum Jang** Methodology, Investigation **Ruth Nussinov** Conceptualization, Methodology, Project administration, Funding acquisition, Resources, Supervision, Writing – original draft & editing **Mardo Kõivomägi** Conceptualization, Methodology, Project administration, Funding acquisition, Resources, Supervision, Visualization, Writing – original draft & editing.

