## Supplementary Information for "Distinct allosteric networks in CDK4 and CDK6 in the cell cycle and in drug resistance"

[4]Department of Human Molecular Genetics and Biochemistry, Sackler School of Medicine, Tel Aviv University, Tel Aviv 69978, Israel

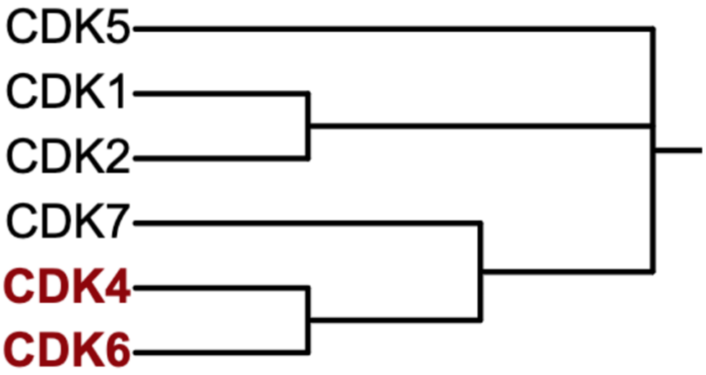


**Figure S1. Phylogenetic tree of human cell cycle CDKs.** It shows the evolutionary closeness of CDK4 and CDK6, compared to the rest of the cell cycle CDKs. The figure is visualized using [1].

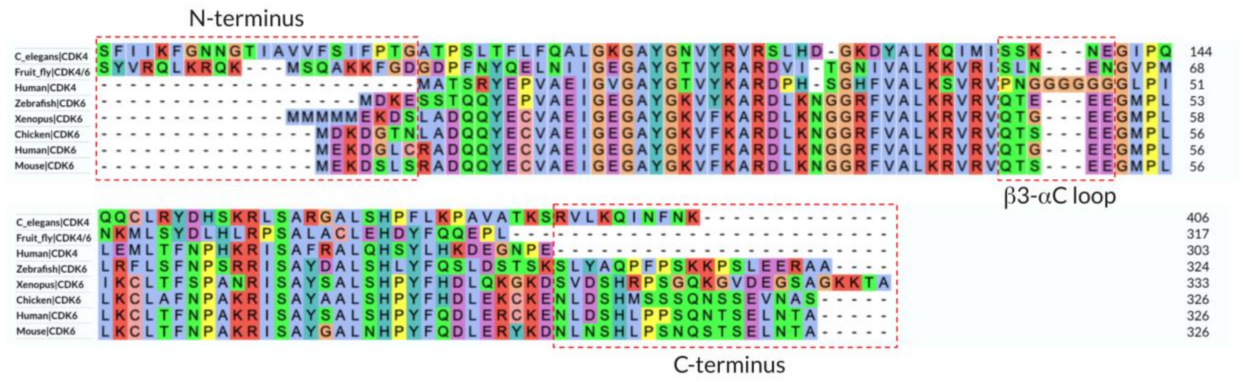


**Figure S2. Multi-species sequence alignment comparing the N- and C-terminal region and β3-αC loop across CDK 4 and CDK6**. The sequence divergence between human CDK4 and CDK6 is mostly conserved in other species as well. The alignment is colored by amino acid properties (red = positive charge, magenta= negative charge, blue = hydrophobic, green = polar, pink = cysteines, orange = glycines, yellow = prolines, cyan = aromatic) to highlight sequence conservation and variation. The key regions (N- and C-termini, and β3-αC loop) are highlighted in red boxes. Each row represents a CDK4 or CDK6 of species ortholog.

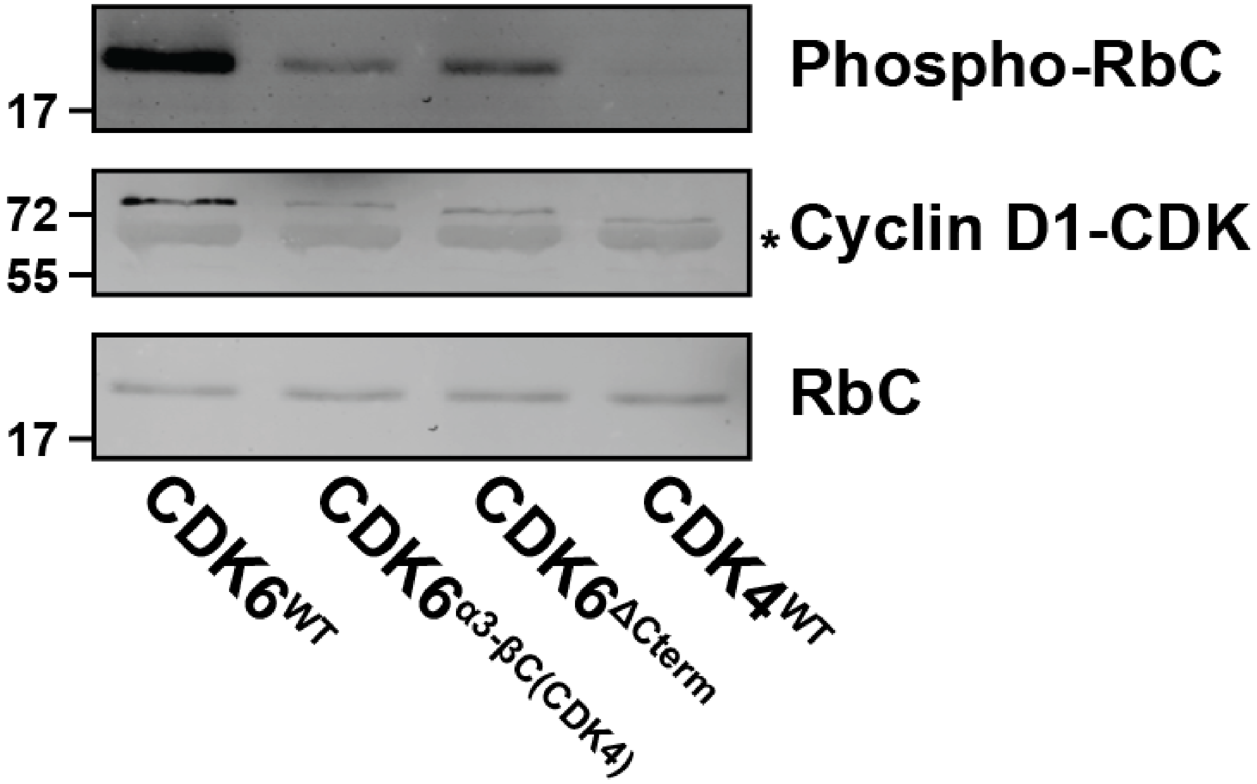


**Figure S3: Representative immunoblot of RbC phosphorylation by Cyclin D1-CDK4/6 complexes.** *In vitro* phosphorylation of RbC by CDK6 (WT, β3-αC(CDK4), and ΔCterm) and CDK4 (WT). Shown is phosphorylated RbC (detected by anti-thiophosphate ester antibodies), cyclin D1-CDK fusions (detected by anti-FLAG antibodies), and RbC detected by total protein stain.

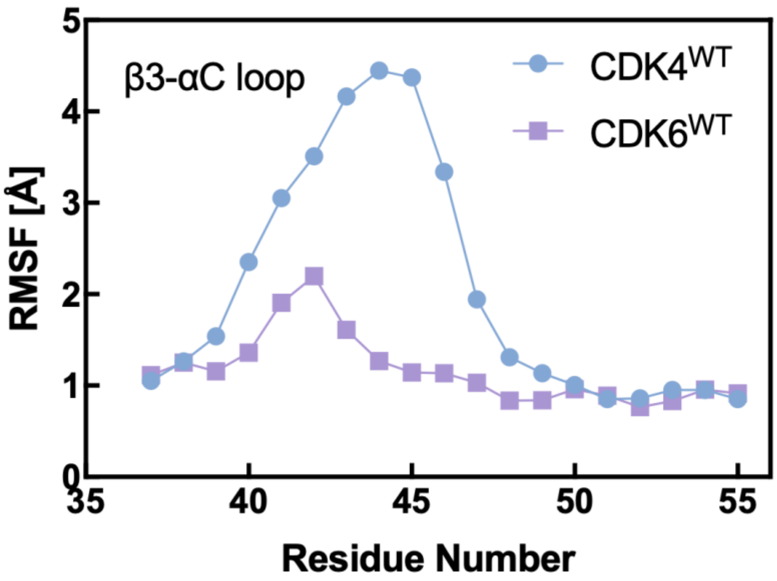


**Figure S4. Root mean square fluctuations (RMSFs) of β3-αC loop in CDK4$^{WT}$ and CDK6$^{WT}$.** The β3-αC loop in CDK4$^{WT}$ has significantly more fluctuation than that in CDK6$^{WT}$. Residue number for CDK6 is subtracted by 8 to match the β3-αC loop residues in CDK4.

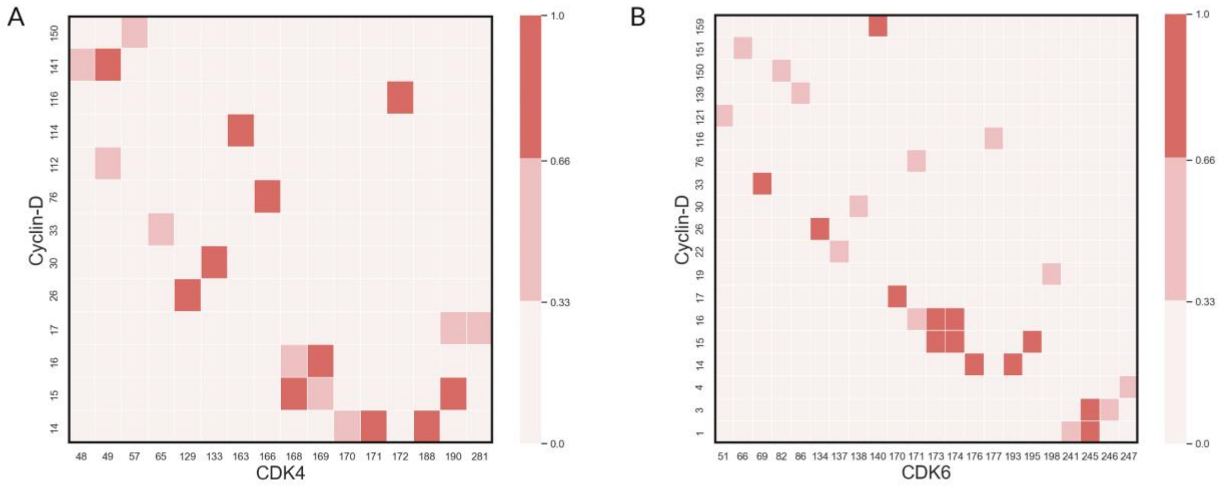


**Figure S5. Contacting residue map between CDK and cyclin for (A) cyclin-D/CDK4, and (B) cyclin-D/CDK6 complexes.** The x-axis represents residue numbers from CDK4/6, while the y-axis shows cyclin-D residues. The color intensity indicates contact frequency between residue pairs during the simulation, with darker red representing higher frequency contacts (scale from 0.0 to 1.0). These maps show more extensive contacting residues between CDK6 and cyclin-D, resulting in its stronger binding affinity.

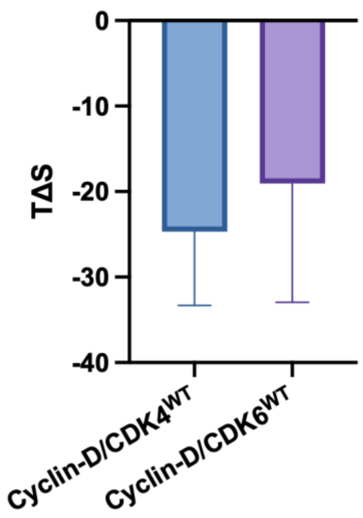


**Figure S6. Entropic contribution to the binding free energy between cyclin-D and either CDK4$^{WT}$ or CDK6$^{WT}$**. The higher degree of entropic contribution of CDK6 complex decreases its binding free energy with cyclin-D.

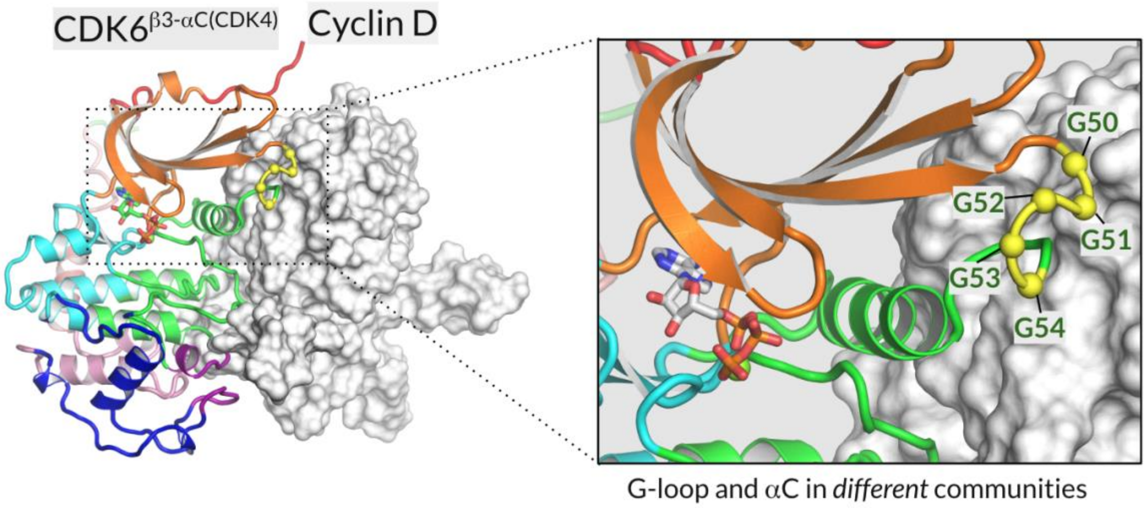


**Figure S7. Community analysis of CDK6$^{WT}$ and CDK6$^{β3\text{-}αC(CDK4)}$.** It shows the impact of the longer CDK4 β3-αC loop can disrupt community organization breaking the coupling between β-strands in C-lobe and αC-helix in CDK6$^{WT}$ (**Fig. 3**). CDK6 is shown in cartoon representation with different colors representing distinct dynamic communities identified through community network analysis. Cyclin-D is displayed in surface representation (gray), highlighting the protein-protein interface. In the zoomed-in view of the G-loop and αC-helix, the G-loop residues (G50-G54) are highlighted in yellow against the cyclin-D surface (gray). The separation of the G-loop and αC-helix into different dynamic communities (shown by distinct colors: orange and green) demonstrates how incorporating the CDK4 β3-αC loop disrupts the normal coupling between these regulatory elements.

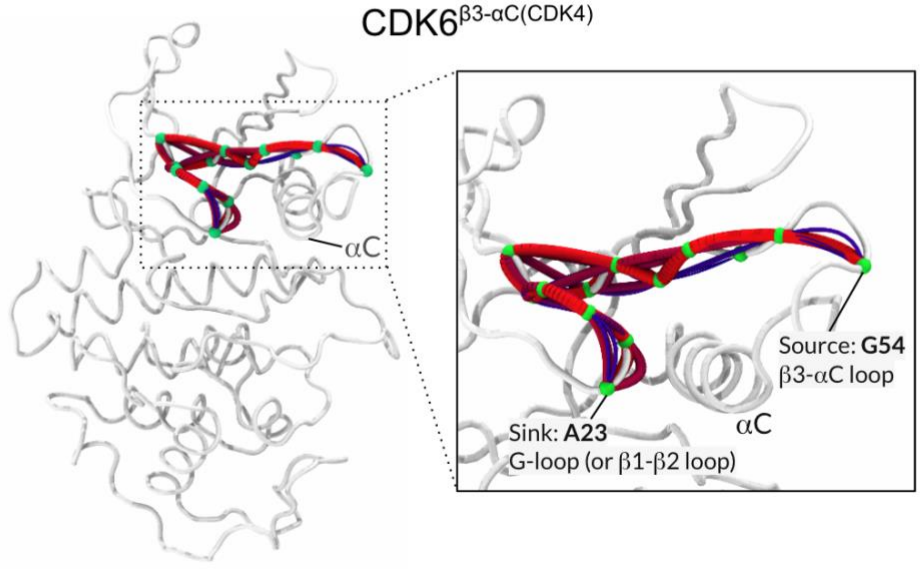


| | | |
|---|---|---|
| CDK4 | VGAYGTVYKA RDPH-SGHFV ALKSVRVPNG GGGGGGLPIS | 52 |
| CDK6 | EGAYGKVFKA RDLKNGGRFV ALKRVRVQTG ---EEGMPLS | 57 |
| $CDK6^{\beta3\text{-}\alpha C(CDK4)}$ | EGAYGKVFKA RDLKNGGRFV ALKRVRVPNG GGGGGGMPLS | 60 |
| | G-loop β3-αC loop | |

**Figure S8. Impact of β3-αC loop substitution on allosteric communication in $CDK6^{\beta3\text{-}\alpha C(CDK4)}$**. Structural visualization of allosteric pathways in $CDK6^{\beta3\text{-}\alpha C(CDK4)}$. $CDK6^{\beta3\text{-}\alpha C(CDK4)}$ is mutant of CDK6 with β3-αC loop swapped for that from CDK4. The zoomed-in view of the allosteric pathways between source residue A29 (in β3-αC loop) and G54 in the G-loop. It exhibits similar allosteric pathways compared to that of CDK4 in **Fig. 4**. Swapping β3-αC weaken the allosteric network between the β3-αC and G-loop, resulting in more dynamic G-loop (β1-β2 loop).

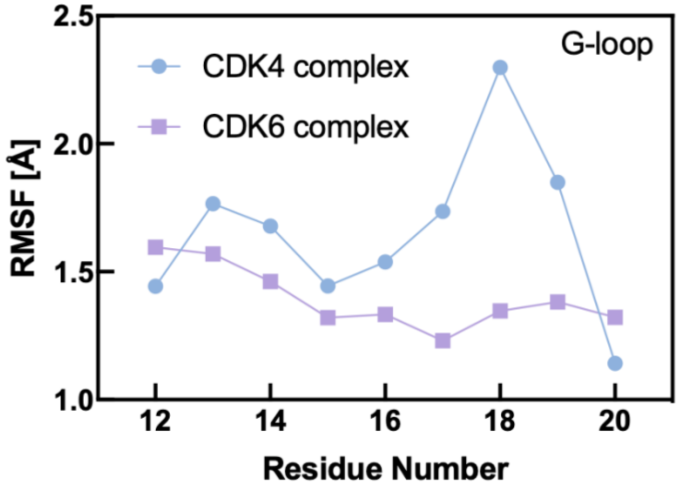


**Figure S9. Root mean square fluctuations (RMSFs) of G-loop in CDK4 and CDK6.** The G-loop of CDK4 has significantly more fluctuation than that in CDK6. The residue number for CDK6 is subtracted by 7 to match the G-loop residues in CDK4.

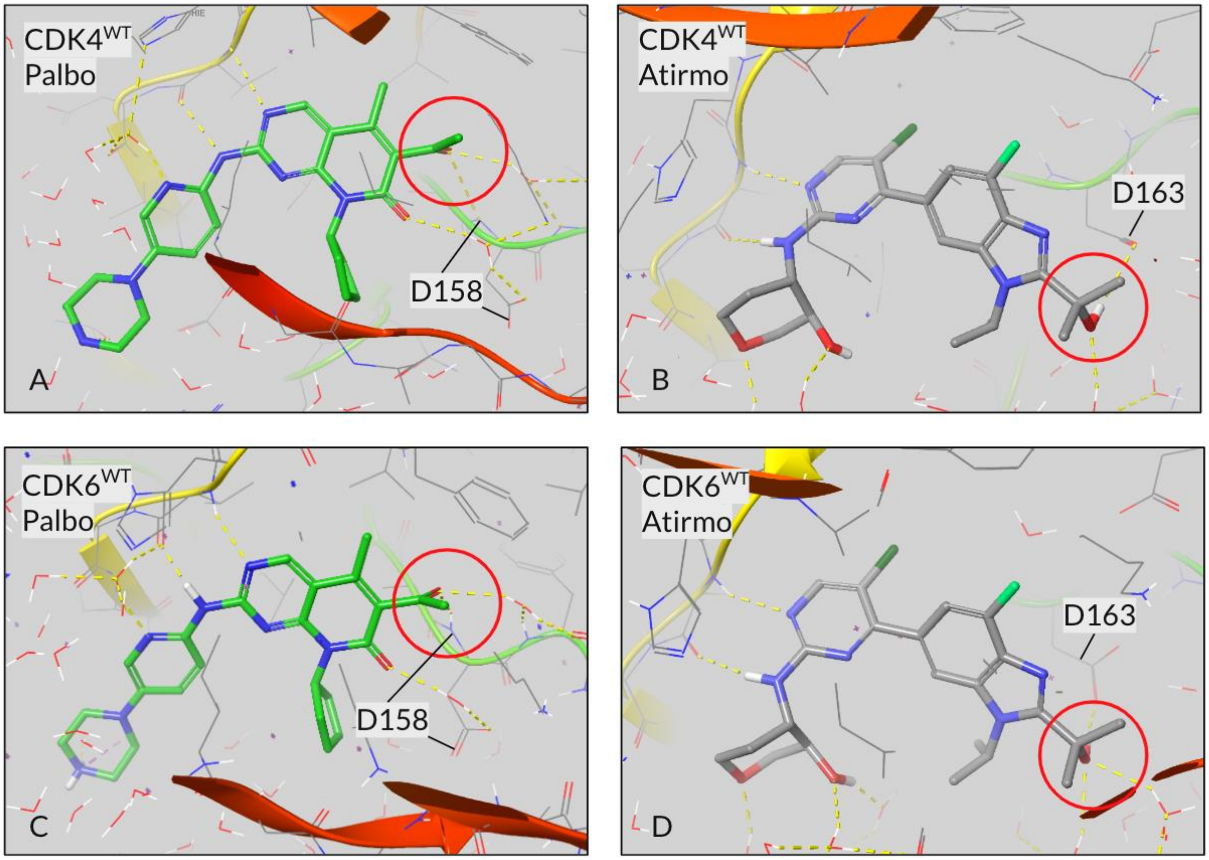


**Figure S10. Comparison of binding modes of palbociclib (PD 0332991) and atirmociclib (PF-07220060) in CDK4 and CDK6 active sites.** Both inhibitors interaction with the aspartic acid (D158 in CDK4, D163 in CDK6) in the DFG motif. (A) CDK4-palbociclib. (B) CDK4-atirmociclib. (C) CDK6-palbociclib. (D) CDK6-atirmociclib. Red circles highlight the carboxylic acid in palbocicilib and isopropyl alcohol in atirmociclib. The additional methyl group in isopropyl alcohol of atirmociclib is responsible for the selectivity of CDK4 due to the flexible G-loop right above this allowing for this bulkier group, in contrast to the less flexible G-loop in CDK6.

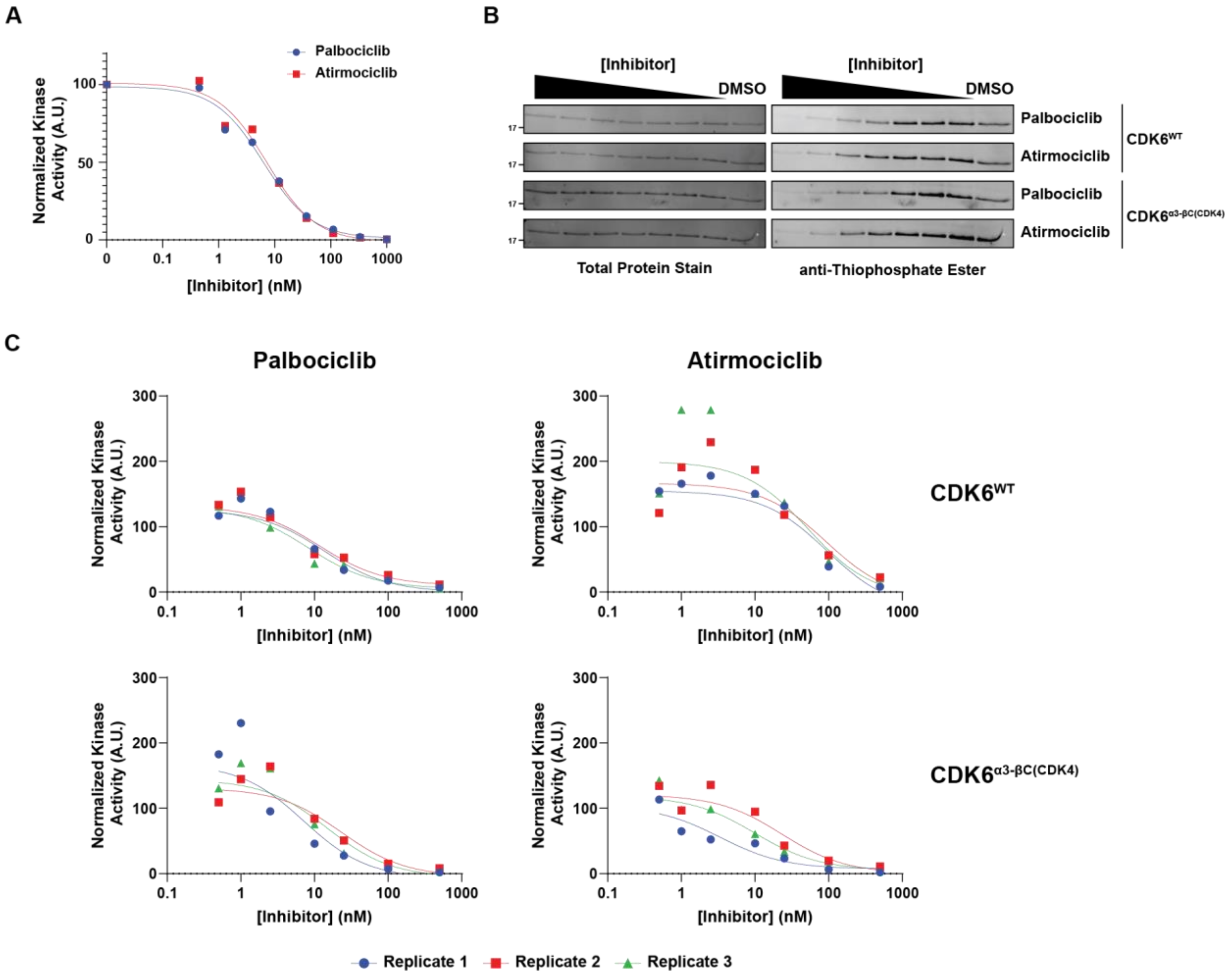


**Figure S11.** ***In vitro*** **CDK4/6 inhibition by palbociclib and atirmociclib.** (A) Palbociclib and atirmociclib mediated inhibition of CDK4 kinase activity toward RbC. (B) Representative immunoblots of palbociclib and atirmociclib mediated inhibition of $CDK6^{WT}$ and $CDK6^{\beta3\text{-}\alpha C(CDK4)}$ kinase activity toward RbC. (C) Individual inhibition curves used to calculate $IC_{50}$ values displayed in Fig. 4I.

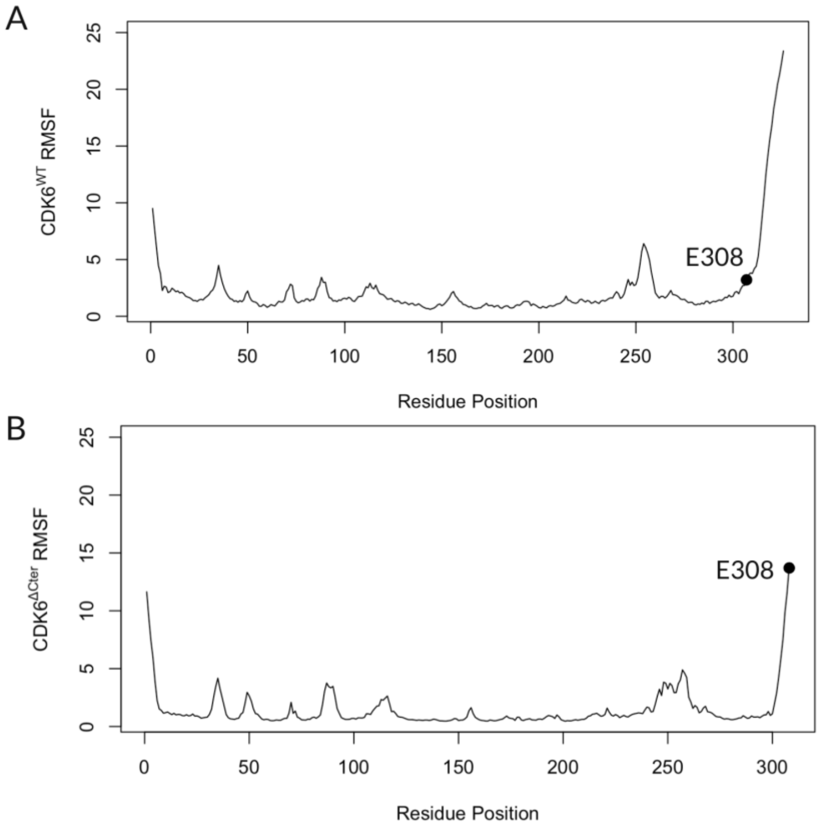


**Figure S12. Root mean square fluctuations (RMSFs) of CDK6$^{WT}$ and CDK6$^{\Delta Cterm}$.** E308, the last residue in CDK6$^{\Delta Cterm}$ has significantly more fluctuation than the same residue in CDK6$^{WT}$. The truncation of C-terminus results in an enhanced fluctuation in the last a few residues in the C-lobe, which leads to the disruption of the intrinsics dynamics of the kinase. For CDK6$^{WT}$, the longer C-terminus shielded E308 from significant fluctuation.

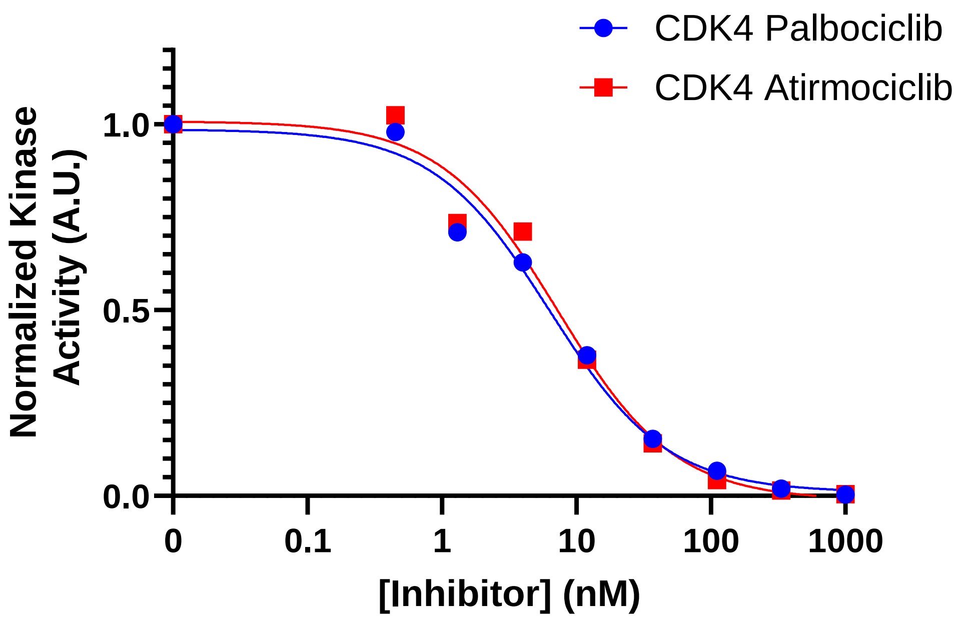


**Figure S13. Dose-response curves showing the effect of CDK4/6 inhibitors, palbociclib (PD 0332991) and atirmociclib (PF-07220060), on the phosphorylation of the retinoblastoma protein (pRbC) in CDK4 complexes.** The x-axis is the concentration of the inhibitors on a logarithmic scale, and the y-axis shows normalized pRb phosphorylation levels. Each curve represents a different condition: CDK4 with palbociclib (blue circles), CDK4 with atirmociclib (red squares).

**Table S1. Summary of the simulations of CDK4 and CDK6 complexes. The mutations of CDKs are shown in Table S2.**

| System | Label | Source PDB | Components | αC | A-loop | DFG | Simulation time (μs) | Number of replicas |
|---|---|---|---|---|---|---|---|---|
| Cyclin-D1/CDK4 (pT172) | $CDK4^{WT}$ | 7SJ3, 2W9Z | Cyclin-D1, CDK4 (pT172), ATP, $Mg^{2+}$ | in | extended | in | 2 | 3 |
| Cyclin-D1/CDK6 (pT177) | $CDK6^{WT}$ | 1XO2, 2W9Z | Cyclin-D1, CDK6 (pT177), ATP, $Mg^{2+}$ | in | extended | in | 2 | 3 |
| Cyclin-D1/$CDK6^{\Delta Cterm}$ (pT177) | $CDK6^{\Delta Cterm}$ | 1XO2, 2W9Z | Cyclin-D1, CDK6 (pT177), ATP, $Mg^{2+}$ | in | extended | in | 2 | 3 |
| Cyclin-D1/$CDK4^{\beta3\text{-}\alpha C(CDK6)}$ (pT172) | $CDK4^{\beta3\text{-}\alpha C(CDK6)}$ | 7SJ3, 2W9Z | Cyclin-D1, $CDK4^{\beta3\text{-}\alpha C(CDK6)}$ (pT172), ATP, $Mg^{2+}$ | in | extended | in | 2 | 3 |
| Cyclin-D1/$CDK6^{\beta3\text{-}\alpha C(CDK4)}$ (pT177) | $CDK6^{\beta3\text{-}\alpha C(CDK4)}$ | 1XO2, 2W9Z | Cyclin-D1, $CDK6^{\beta3\text{-}\alpha C(CDK4)}$ (pT177), ATP, $Mg^{2+}$ | in | extended | in | 2 | 3 |

**Table S2. CDK4 and CDK6 and mutant sequences.** Altered sequences are underlined; delete sequence is denoted as red dash.

| Name | Sequence | Ending residue ID |
|---|---|---|
| **$CDK4^{WT}$** | -------MAT SRYEPVAEIG VGAYGTVYKA RDPH-SGHFV ALKSVRVPNG GGGGGGLPIS | 52 |
| **$CDK6^{WT}$** | MEKDGLCRAD QQYECVAEIG EGAYGKVFKA RDLKNGGRFV ALKRVRVQTG ---EEGMPLS | 57 |
| **$CDK6^{\beta3\text{-}\alpha C(CDK4)}$** | MEKDGLCRAD QQYECVAEIG EGAYGKVFKA RDLKNGGRFV ALKRVRVPNG GGGGGGMPLS | 60 |
| **$CDK4^{\beta3\text{-}\alpha C(CDK6)}$** | -------MAT SRYEPVAEIG VGAYGTVYKA RDPH-SGHFV ALKSVRVQTG ---EEGLPIS | 57 |
| **$CDK6^{\Delta Cterm}$** | SALSHPYFQD LERCKE------------------ | 308 |

**Table S3: Yeast strains used in this study.**

| Strain | Protein Expressed | Mutation |
|---|---|---|
| pBTB_003-1 | pRS425-pGAL1-3XFLAG-CCND1-Linker-CDK4 | Wild Type |

| | | |
|---|---|---|
| pBTB_004-1 | pRS425-pGAL1-3XFLAG-CCND1-Linker-CDK6 | Wild Type |
| MK_1149 | pRS425-pGAL1-3XFLAG-CCND1-Linker-CDK6 | β3-αC(CDK4) |
| yBT73 | pRS425-pLEXA-3XFLAG-CCND1-Linker-Cdk6 | ΔCterm |
